# Analysis of behavioral flow resolves latent phenotypes

**DOI:** 10.1101/2023.07.27.550778

**Authors:** Lukas M. von Ziegler, Fabienne K. Roessler, Oliver Sturman, Rebecca Waag, Mattia Privitera, Sian N Duss, Eoin C. O’Connor, Johannes Bohacek

## Abstract

The nuanced detection of rodent behavior in preclinical biomedical research is essential for understanding disease conditions, genetic phenotypes, and internal states. Recent advances in machine vision and artificial intelligence have popularized data-driven methods that segment complex animal behavior into clusters of behavioral motifs. However, despite the rapid progress, several challenges remain: Statistical power typically decreases due to multiple testing correction, poor transferability of clustering approaches across experiments limits practical applications, and individual differences in behavior are not considered. Here, we introduce “behavioral flow analysis” (BFA), which creates a single metric for all observed transitions between behavioral motifs. Then, we establish a “classifier-in-the-middle” approach to stabilize clusters and enable transferability of our analyses across datasets. Finally, we combine these approaches with dimensionality reduction techniques, enabling “behavioral flow fingerprinting” (BFF) for individual animal assessment. We validate our approaches across large behavioral datasets with a total of 443 open field recordings that we make publicly available, comparing various stress protocols with pharmacologic and brain-circuit interventions. Our analysis pipeline is compatible with a range of established clustering approaches, it increases statistical power compared to conventional techniques, and has strong reproducibility across experiments within and across laboratories. The efficient individual phenotyping allows us to classify stress-responsiveness and predict future behavior. This approach aligns with animal welfare regulations by reducing animal numbers, and enhancing information extracted from experimental animals

## Introduction

The majority of preclinical biomedical research is conducted in mice. Thus, the reliable detection of complex mouse behavior is critical to gain insights into disease conditions, genetic phenotypes and internal states (emotion, well-being etc.). Traditionally, behavior assessment has been performed by manual observation of animals^1–3^, a time consuming approach notoriously plagued by human bias, high inter- and intra-rater variability and poor reproducibility^4, 5^. The advent of pose-estimation technology^6–8^ has enabled the automation of this process using supervised machine learning methods, thus solving the problem of variability and reproducibility^5, 9–13^. However, the process of selecting and defining the relevant behaviors still introduces considerable human bias and limits the analyses to predefined assumptions about animal behavior^4, 14, 15^.

Given these limitations, data-driven methods that analyze behavior independent of human intervention have recently attracted more interest. Based on raw or processed video data, these approaches segment behavior using clustering algorithms or more sophisticated state-space models, and have revealed a remarkable complexity underlying even the simplest behavior tasks^16–21^. For example, open field behavior - routinely used as a testbed for analyzing unconstrained rodent behavior - was shown to contain a large number of readily identifiable behavioral clusters (also called syllables or motifs)^16, 18, 19, 21^.

Despite this rapid progress, such advancements have given rise to three novel challenges, which we address in this work. 1) The multiple testing problem: One of the major promises of unbiased, data-driven analyses of animal behavior is the ability to better resolve differences between experimental groups, by analyzing large numbers of behavioral variables^4, 14, 15^. However, the more variables we identify, the less power we have to detect group differences due to increased statistical demands from multiple testing corrections. We solve this by introducing a “behavioral flow analysis” (BFA), which uses a single metric to identify treatment effects based on the observed transitions between the different behavioral clusters. 2) The transferability problem: Clusters that represent behaviors vary between experiments, making it impossible to compare clustering results between different experiments. Further, it remains difficult to transfer a model between different testing conditions, datasets or laboratories. We address this using a “classifier-in-the-middle” approach, which trains a supervised machine learning classifier to recognize established behavioral clusters in newly encountered datasets. 3) The problem of individual differences: Despite ever more nuanced analyses on the group level, it has not yet been demonstrated that behavioral personality profiling - akin to a detailed clinical assessment routinely conducted in human studies - is possible in mice. We combine BFA and stabilized clusters with dimensionality reduction techniques to generate a single, high-dimensional datapoint for each animal. This “behavioral flow fingerprinting” (BFF) approach provides an individual assessment of each animal and allows large-scale comparisons of animal behavior across a wide range of experimental manipulations.

Using a series of stress paradigms, pharmacological and circuit neuroscience interventions, we test and validate our approach across many behavioral datasets. Our results are in line with the reduce-and-refine principles set forth by animal welfare regulations, as our approach increases statistical power, reduces the number of animals required for experiments, and increases the information extracted from each experimental animal.

## Results

### 1. Behavior Flow Analysis (BFA) increases power to detect treatment effects

#### Data-driven clustering vs. classical analyses

The ability to resolve nuanced behaviors in a data-driven manner holds the promise to reliably and powerfully identify behavioral phenotypes^16–21^. We hypothesized that such an unsupervised approach would reveal known phenotypes in existing datasets with greater sensitivity. To test this, we first turned to a large, published behavioral dataset^22^ in which mice were exposed to chronic social instability stress (CSI, n=30) or to control handling (n=29), before being tested on the open field test (OFT) for 10 minutes (Figure 1A). For classical behavior analysis - as described previously^9, 22^ - we tracked 13 bodypoints using the pose-estimation tool DeepLabCut^6, 7^, transformed the point-tracking data into a set of features, computed standard OFT parameters (’distance traveled’ and ’time in center’), and employed a supervised machine learning analysis to quantify supported and unsupported rears. CSI mice spent more time in the center of the open field and they traveled more distance, while their rearing behavior was not affected (Figure 1B), thus representing a useful dataset with known phenotypical differences between groups.

**Figure 1.**
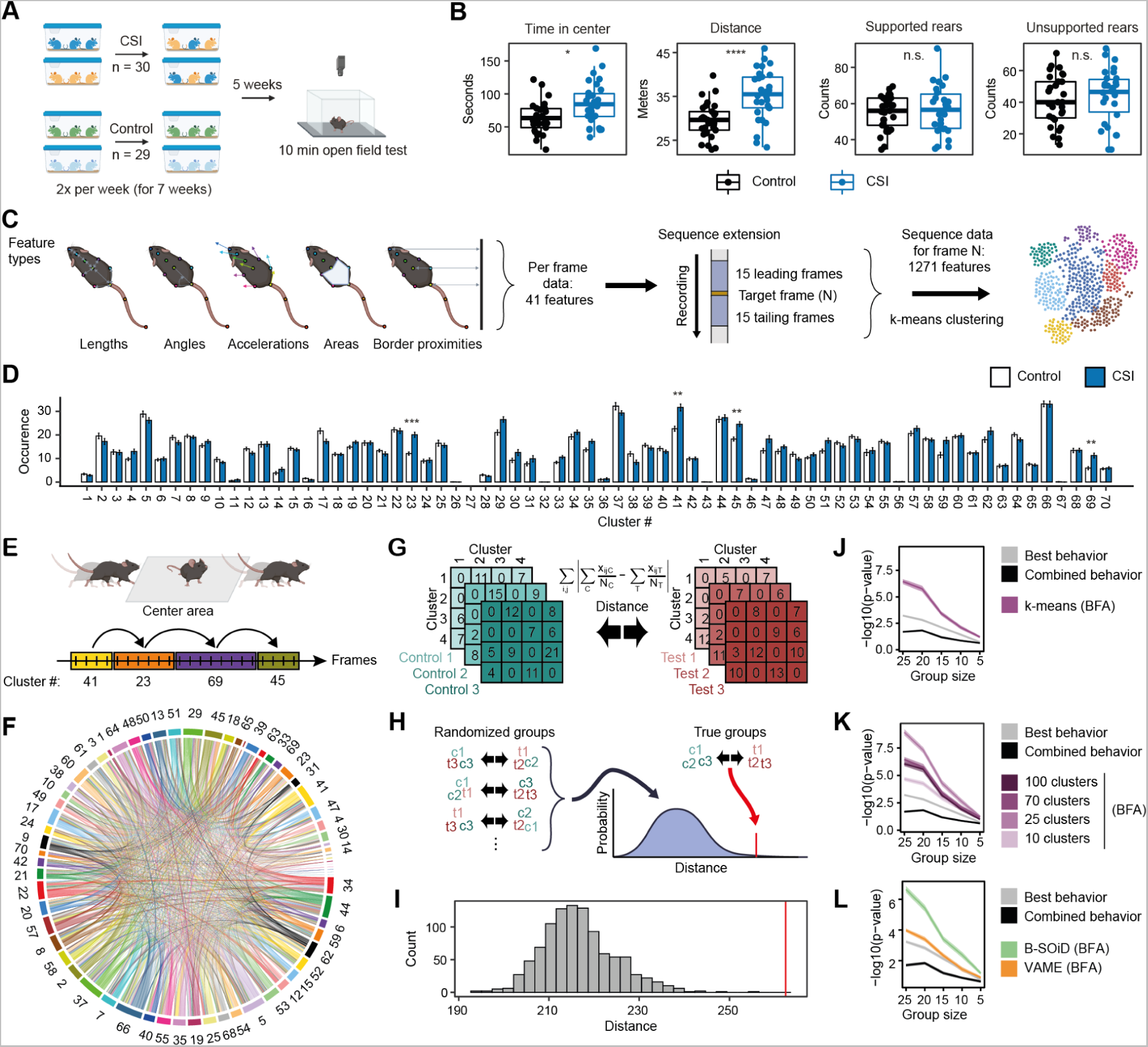
Behavioral flow analysis increases sensitivity to detect phenotypes. (A) Schematic showing experimental design for chronic social instability (CSI). (B) Classical behavior readouts in the open field test show that CSI mice spend more time in the center (t(53)=2.96, adj. p=0.045) and travel greater distance (t(52)=4.55, adj. p<0.001). (C) Feature extraction based on pose-estimation tracking and sequential feature integration for subsequent clustering. (D) k-means cluster occurrence in CSI. (E) Schematic example of behavioral flow based on cluster transitions. (F) Average behavioral flow over all animals. (G) Schematic of computing Manhattan distance to compare behavioral transition matrices between groups. (H) Bootstrapping approach used for behavioral flow analysis (BFA) to compare the transition distance based on the true group assignment versus the randomized group assignment. (I) BFA reveals a treatment effect for CSI (percentile=99.9, z=5.72, p<0.001). (J) BFA enhances the sensitivity to detect the CSI phenotype. (K) BFA enhances sensitivity using various numbers of k-means clusters. (L) (J) Phenotype detection sensitivity in CSI using B-SOiD or VAME clustering with BFA. Adj. p-values are denoted as: *<0.05, **<0.01, ***<0.001. Error bars denote ± SEM.

We then tested how well an unsupervised clustering approach performs at resolving group differences. We first transformed the point-tracking data into an extended set of 41 features (see Methods section). These features were then resolved over a sliding time window (+/-15 frames) to describe short temporally resolved sequences (Figure 1C). The resulting data contained 1271 dimensions for each frame. We then used a simple and computationally efficient k-means clustering algorithm to resolve different behaviors. To determine the best number of clusters, we chose an approach used previously for segmenting behavior by VAME^21^ and MoSeq^17^. To this end, we first partitioned the recorded behavior into 100 clusters, and then chose the number of clusters which represented 95% of the imaging frames. This mark indicated about 70 clusters (Suppl. Figure 1A), so we subsequently re-ran the clustering approach using 70 clusters. Although nominal p-values revealed that CSI and control animals behaved differently on many of these clusters, only 4 out of 70 clusters survived the appropriate multiple testing correction (Benjamini-Yekutieli, Figure 1D). Visual inspection of these significant clusters reveals - in agreement with the classical analysis - that they capture the time the animals spent in the center of the open field. Specifically, these four clusters represent movement of the mouse from the periphery into the center (cluster 41, Suppl. Video 1), movement in or through the center (cluster 23, Suppl. Video 2), orienting and turning in the center (cluster 69, Suppl. Video 3) and movement from the center back to the periphery (cluster 45, Suppl. Video 4).

#### Quantifying transitions between clusters

Importantly, behavior emerges as a dynamic property of moment-to-moment action. Thus, we took advantage of the frame-by-frame resolution of the clustering data to assess the behavioral flow (i.e. the temporal sequence in which one cluster transitions to another cluster) in each animal (Figure 1E). To enable this approach, we created an open source code base that allows the loading and analysis of (manual, supervised or unsupervised) labeling results on a frame-by-frame basis. We used this package to first analyze how behavior flows across all animals in the CSI experiment, independent of group assignment (Figure 1F). This demonstrated that many clusters had ingoing and outgoing transitions that were much more likely to occur. An example are the clusters identified as significant between CSI and control mice, where behavioral flow indicates that when mice move from the periphery to the center (cluster 41) this can be followed by movement through the center (cluster 23) or exploration in the center (cluster 69), which is most often followed by a movement back into the periphery (this “crossing of the open field”- sequence is displayed schematically in Figure 1E, and supported by data in Figure 1F).

We then asked whether these transitions would reveal group differences between CSI and control mice. However, when we applied multiple testing corrections to account for the high number of observed transitions (1753 observed transitions out of 4830 possible transitions), none of them remained significant (adj. p>0.05). To demonstrate that the information gain is punished by multiple testing correction, we designed a sensitivity assay. We used the CSI dataset to generate multiple two-group comparisons of random subsets of each group of mice, and successively reduced group sizes *in silico* (Suppl. Figure 1B). The results demonstrate that - when using unadjusted p-values - both cluster usage and transition occurrences perform better than classical analyses (for which the best statistical value between distance, time in center, supported rearing and unsupported rearing was used; Suppl. Figure 1C). However, when applying multiple testing corrections, both cluster and transition measures perform poorer than classical analyses in detecting phenotype differences (Suppl. Figure 1D).

#### Behavioral Flow Analysis (BFA)

To address the multiple testing problem, we used a statistical approach to detect group differences based on the combined behavioral flow data, which we term “behavioral flow analysis” (BFA). The BFA method first defines the difference between the two experimental groups based on the Manhattan distance between group means across all behavioral transitions (Figure 1G). To assess if this distance is significantly larger than expected, we used a bootstrapping approach where randomized group assignments were generated using the original data to estimate a null distribution of the inter-group distance (Figure 1H). Then, we calculated the percentile and tested the true distance against the null distribution using a right-tailed z-test, which revealed a strong group difference that is very unlikely due to chance (Figure 1I). Given this result, we tested whether BFA would increase the power to detect differences between groups. We again used the sensitivity assay by gradually lowering group sizes from 25 to 5 animals *in silico*. With large group numbers (>10mice/group), the increase in statistical power provided by BFA is extremely large (Figure 1J). Gradually lowering the group sizes from 25 to 5 replicates demonstrated that BFA still resolved a phenotype at lower group sizes than the best of the classically recorded behaviors (termed “best behavior”, see Methods for details, Figure 1J).

To ensure that this is not due to potentially poor manual selection of relevant behaviors we devised a combined behavior analysis, which leverages all data generated with the OFT analysis using DLC (as described in a previous publication^9^, DLCAnalyzer) in combination with a principal component analysis (PCA) to deal with the highly correlated variables. We observed that this composite measure (termed “combined behavior”) performed worse than the “best behavior” (Figure 1J), confirming that by manually selecting the “best behavior” we created an advantageous scenario that requires prior knowledge of the phenotype. Thus, BFA is indeed much more powerful at detecting group differences than the behavior that best captures group differences, and more powerful than a composite score of predictive individual behaviors.

#### BFA works with various clustering approaches

As many different clustering approaches have been successfully used to analyze rodent behavior^16, 19, 21^, we tested whether the BFA method would work independent of the clustering algorithm employed. We first turned to VAME, one of the most recently developed unsupervised approaches for behavior analysis. DLC-based tracking data was used for egocentric alignment as described by the authors (Suppl. Figure 2A) and reached good model performance (Suppl. Figure 2B). Estimating the best number of clusters (as described above) we selected 80 clusters (Suppl. Figure 2C). Similar to the k-means approach, 5 of these clusters revealed significant group differences (adj. p<0.05) between CSI and control animals (Suppl. Figure 2D), despite the fact that both algorithms only showed moderate alignment in the type of clusters identified (Suppl. Figure 2E). The resulting behavioral flow diagram showed differences for certain transitions (Suppl. Figure 2F), but the attempt to analyze all possible behavioral transitions was again punished by multiple testing correction (1820 observed transitions out of 6320 possible transitions), revealing no statistically significant group differences (Suppl. Figure 2G). However, BFA - computed based on all behavioral transitions - was able to reveal a highly significant group difference (Suppl. Figure 2H).

#### BFA works with various clustering approaches and cluster numbers

We also tested BFA using clusters generated by B-SOiD, another popular unsupervised clustering pipeline^16^. Using B-SOiD restricted the input features to 9 body points (Suppl. Figure 3A). Adjusting the number of clusters in B-SOiD, which uses a hierarchical clustering method, is not straight-forward, and systematically varying the input parameters generated a wide range of possible clusters. We settled for 8 clusters, which efficiently separated the data (Suppl. Figure 3B), and which were accurately assigned to single frames by the random forest classifier as assessed by confusion matrix and test data accuracy (Suppl. Figure 3C). It has indeed been reported recently that some clustering algorithms only resolve a few behavior motifs in open field data^19^, and it offered an opportunity to test whether our BFA approach would work on fewer clusters. Three of the identified clusters showed significant group differences (adj. p<0.05; Suppl. Figure 3D). Mapping these B-SOiD clusters to our k-means clusters reveals that every B-SOiD cluster contains many of the clusters represented by k-means (Supp. Figure 3E). The resulting behavioral flow diagrams show that 5 transitions were significantly different between CSI and controls (adj. p<0.05, Suppl. Figure 3 F-H), in line with the notion that the small number of observed transitions (56 observed transitions out of 56 possible transitions) rendered the multiple testing correction less punishing. We then conducted BFA and again found that it was very powerful in detecting a group difference (Suppl. Figure 3I). A systematic assessment showed that BFA can increase statistical power across all three clustering approaches (Figure 1J/L). B-SOiD yielded good results despite the small number of clusters, thus we also compared the performance of BFA using the k-means-based approach with different numbers of clusters. Indeed, a broad range of cluster numbers (from 10 to 100) all strongly increased statistical power (Figure 1K). Because 25 clusters showed the best results in this dataset, we used 25 clusters in the subsequent analyses. Overall, BFA enables efficient phenotype detection across clustering algorithms, and even with very low cluster numbers.

### 2. Cluster stabilization enables direct comparisons across experiments

#### Adding new behavioral datasets

Because behavior is complex and nuanced, cluster boundaries and sub-divisions tend to be blurry, and as a result unsupervised clustering tends to be sensitive to minor differences in the data. Therefore, some cluster stabilization would be required to compare results across experiments. To address this issue, we added two new behavioral datasets. First, we performed an experiment in which mice were tested 45min and 24hrs after a short swim stress exposure (acute swim (AS), n=15; control, n=15). (Figure 2A). Second, we added a pharmacological stress model that allowed us to introduce a graded stress response by injecting mice with variable doses of yohimbine, an α2-adrenergic receptor antagonist, which triggers strong noradrenaline release by disinhibiting the locus coeruleus^23, 24^. Control animals were injected with saline (n=5), while each mouse in the treatment group received one injection of yohimbine ranging from 0.4 to 6 mg/kg (n=15; Figure 2B).

**Figure 2.**
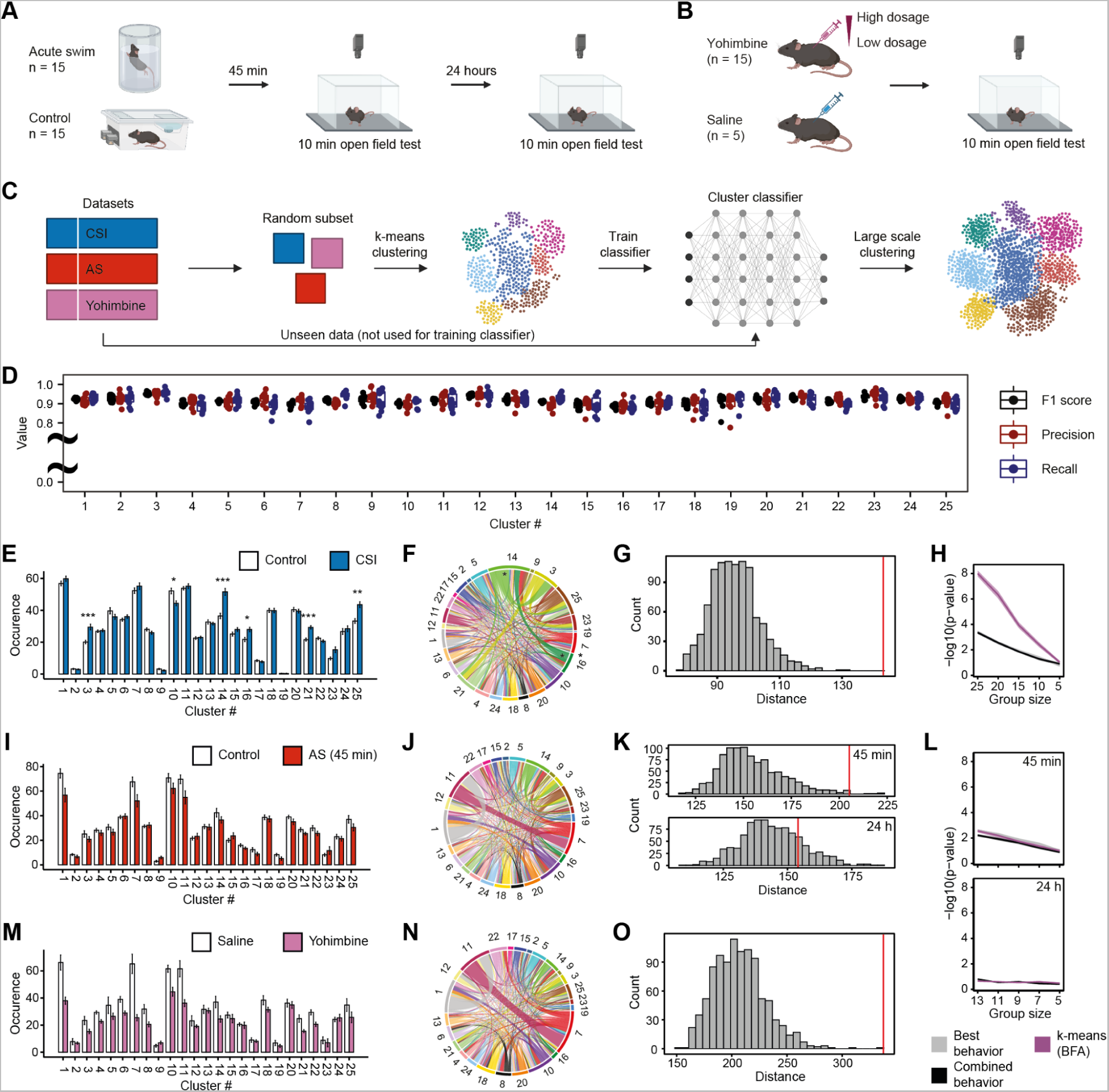
Cluster stabilization enables comparisons across experiments. (A) Schematic of experimental design for OFT after acute swim (AS) stress, or (B) after yohimbine injections. (C) Clustering across large datasets. (D) Classifier performance on 10-fold cross-validation for each cluster. (E) Quantification of cluster occurrences in CSI. (F) Absolute differences in behavioral flow in control vs. CSI. For each cluster, the absolute difference in the observed number of transitions between groups is plotted. (G) BFA reveals a treatment effect for CSI (percentile=99.9, z=6.02, p<0.001). (H) BFA enhances phenotype detection sensitivity in CSI. (I) Cluster occurrences in AS (45 min). (J) Absolute difference in behavioral flow in control vs. AS (45 min). (K) BFA reveals a treatment effect for AS at 45 min (percentile=99.4, z=3.09, p=0.001) but not at 24 h (percentile=80.9, z=0.85, p=0.198). (L) Phenotype detection sensitivity in AS. (M) Cluster occurrences in yohimbine. (N) Absolute difference in behavioral flow in saline vs. yohimbine. (O) BFA reveals a treatment effect for yohimbine (percentile=99.8, z=5.56, p<0.001). Adj. p-values are denoted as: *<0.05, **<0.01, ***<0.001. Error bars denote ± SEM.

#### Cluster stabilization across experiments

To obtain comparable clustering results across these experiments, all the data could be used in one big clustering experiment. However, this would entail re-running the unsupervised clustering every time new data is added and soon become computationally challenging as previously observed^18^. Instead, we stabilized clusters using a classifier-in-the-middle approach. To this end, we first selected only a random set of 20 animals from each behavioral experiment (CSI, AS, yohimbine) for performing k-means clustering to generate 25 clusters, as this number of clusters was sufficient to detect phenotypical differences as shown above (Figure 1K). Afterwards, we used these clustering results to train a neural network that can imitate the clustering on the random subset for the rest of the data (Figure 2C). Using a 10-fold cross validation we found that this approach had good (>0.9) precision and recall values across all 25 clusters (Figure 2D).

Using this cluster stabilization approach, we first re-analyzed the CSI experiment. As we now only used 25 clusters, multiple testing correction was less punishing and we identified 6 significant clusters (Figure 2E), and two significant transitions (Figure 2F). Visual inspection revealed - consistent with the original analysis - that all clusters that occurred more frequently in CSI mice captured behaviors related to active locomotion in the center: movement in the center of the open field (cluster 3, Suppl. Video 5), the initiation of movement from the periphery to the center (cluster 14, Suppl. Video 6), fast locomotion crossing the center or moving from center to periphery (cluster 21, Suppl. Video 7), and exploration/turning in the center or movement towards the periphery (cluster 25, Suppl. Video 8). In contrast, the only underrepresented cluster in CSI mice was movement or exploration along the periphery of the open field (cluster 10, Suppl. Video 9). Accordingly, the transitions significantly overrepresented in CSI mice captured the transition from being stationary close to the wall to initiating movement into the center (cluster 16 to cluster 14), and from moving into the center to slowing down and re-orienting in the center (cluster 14 to cluster 3). Also the BFA pipeline reproduced the very strong phenotype (Figure 2G) and increased statistical power compared to classical analyses (Figure 2H). Taken together, our findings reveal that, independent of the number of clusters, our approach captures meaningful behavioral motifs and interpretable transitions, also when the original dataset is reanalyzed using a set of clusters derived from training on subsets of videos from various experiments.

#### Cluster stabilization across experiments: Swim stress and yohimbine

We then applied the same analysis pipeline using the clustering classifier to open field behavior assessed 45 minutes after an acute swim stress. None of the 25 clusters and none of the observed transitions revealed any group differences (Figure 2I,J), yet BFA readily identified a significant group difference (Figure 2K). When we instead analyzed open field behavior 24hrs after swim stress, neither the 25 clusters or their transitions (Suppl. Figure 4A,B) nor BFA revealed a discernible phenotype anymore (Figure 2K), consistent with our previous observations that swim stress only induces transient changes in mouse behavior^25^. These results suggest that BFA can detect group differences when other methods fail, but it does not create arbitrary effects in a scenario when no effects would be expected. Despite the successful BFA analysis, the sensitivity to detect a phenotype could not be improved in this dataset (Figure 2L).

Next we applied the same analysis pipeline to mice injected with escalating doses of yohimbine. For this, we grouped all yohimbine-injected mice into one group (yohimbine, n=15), and compared them to the 5 saline-injected controls (Figure 2B). Because of the small number of mice in the control group, nominally large effects on various individual clusters did not yield statistical significance after multiple testing correction (Figure 2M), nor on the level of observed transitions (Figure 2N). However, BFA was again able to reveal a highly significant group difference (Figure 2O), showcasing the power of this analysis even when applied to experiments with intentionally high variability and small, otherwise underpowered group sizes.

A visual comparison of cluster occurrence across all three experiments reveals very clearly that acute swim stress and yohimbine injections - which both acutely trigger strong noradrenaline release - induce similar behavioral profiles in the open field, but these behavioral changes are very distinct from the ones induced by CSI (Figure 2E,I,M). For example, yohimbine strongly suppresses clusters 1 and 11, which capture the initiation and termination of a supported rear, respectively (see Suppl. Video 10 and 11), in line with our previous findings that yohimbine reduces supported rearing^24^.

### 3. Behavioral Flow Fingerprinting (BFF) captures individual differences

#### Capturing cluster transitions per animal

Thus far, we have leveraged the power of unsupervised clustering and BFA to identify group differences. Ultimately, however, it is necessary to understand behavior on the level of individual animals. Recently, MoSeq was used to demonstrate that unsupervised approaches can resolve differences between different drugs and drug doses^18^. We thus asked whether behavioral flow would be sensitive to drug dosage in the yohimbine experiment (Figure 3A). When modeling occurrence of transitions as a function of dosage, one transition, from cluster 11 to 7, showed a significant association with the dose of yohimbine (Figure 3B). Visual inspection showed that this transition captured the end of a supported rear (cluster 11, Suppl. Video 11), and the subsequent turning motion and initiation of movement (cluster 7, Suppl. Video 12). At higher doses of yohimbine, mice transition less frequently between these clusters related to supported rearing (Figure 3B). To further explore individual differences based on behavioral flow, we considered that the behavioral flow diagrams represent high-dimensional data for each animal (625 possible transitions between 25 clusters). For easier visualization and analysis, we applied Uniform Manifold Approximation and Projection (UMAP) to project the high-dimensional behavioral transition matrix of each animal onto a single datapoint in 2D space, while retaining as much relevant information as possible. The resulting 2D embeddings showed a very good separation of the saline vs yohimbine groups (Figure 3C), and were also able to resolve low vs high dosages, despite the small animal numbers used in this experiment (Figure 3D). As this high-dimensional representation of behavioral complexity aims to provide a unique description of each animal’s behavioral phenotype, we refer to this approach as “Behavioral Flow Fingerprinting” (BFF).

**Figure 3.**
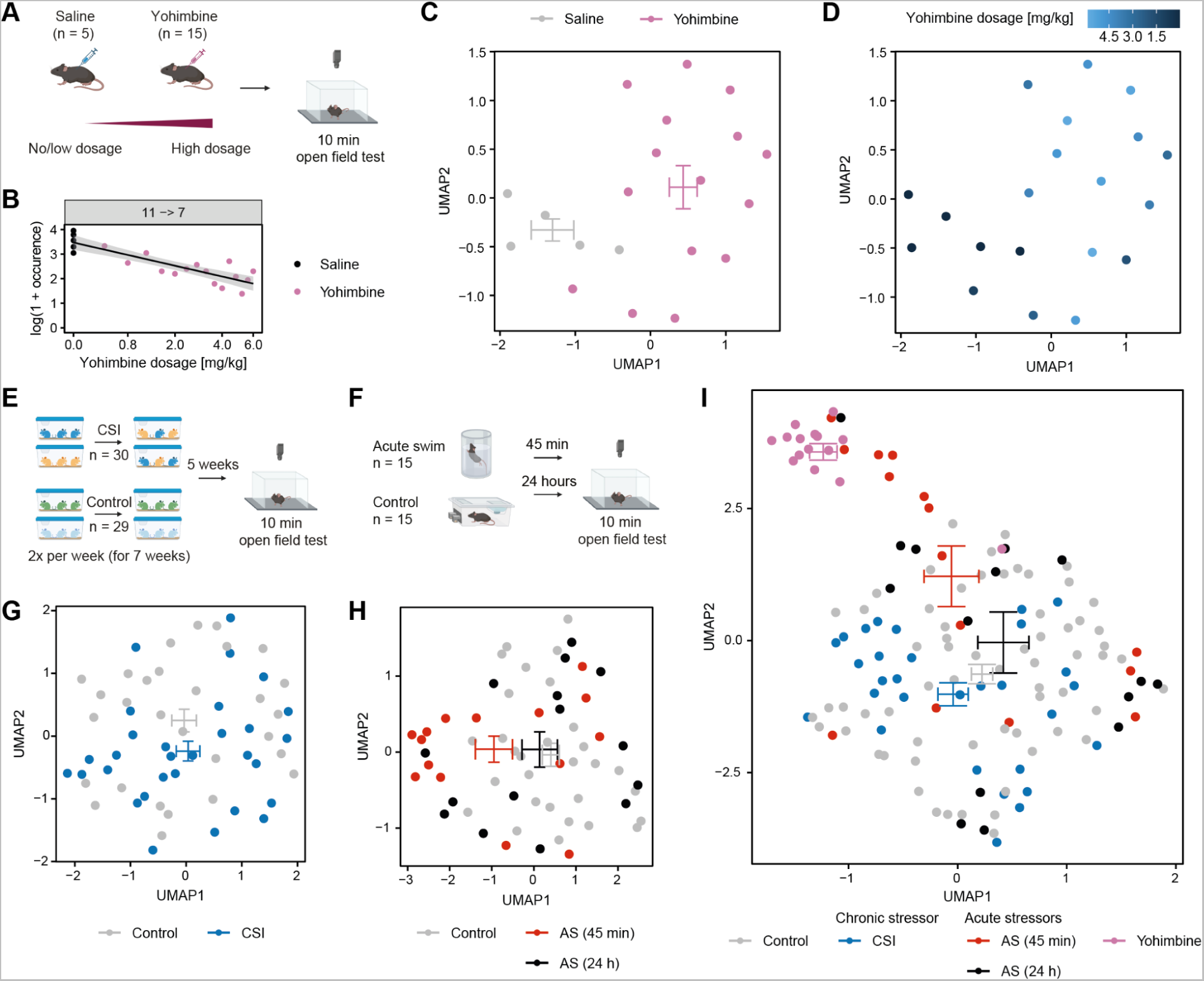
Behavioral flow fingerprinting (BFF) captures individual differences in high-dimensional space. (A) Schematic of experimental design showing the escalating dose of yohimbine. (B) Using dose in a log-linear model reveals one significant transition (R^2^=0.75, F(1,18)=55.44, adj. p=0.002). (C) BFF reveals a separation between mice treated with vehicle versus yohimbine when applying UMAP dimension reduction to their transition matrices. The crossbars represent the average UMAP1 and UMAP2 values with SEM for each group. (D) BFF can also visualize the drug dose delivered to each animal. (E, G) BFF applied to the chronic social instability (CSI) dataset. (F, H) BFF applied to the acute swim stress (AS) dataset. (I) Plotting BFF embeddings across all three experiments (CSI, AS, yohimbine) reveals a separation of all experimental groups in 2D space.

#### Plotting behavioral profiles across many experiments

Being able to stabilize the unsupervised clustering analyses across experiments, and to capture individual behavioral response profiles of each animal, we explored whether we could plot individual behavioral profiles across experiments. We first performed the BFF embedding of all cluster transitions on the CSI dataset, which revealed a shift of CSI animals away from controls (Figure 3E,G). However, it also showed the large overlap between both experimental groups, which reflects the large spread between data points (individual mice) using classical measures such as “distance moved” or “time in center” (see Figure 1B).

We then applied the same BFF analysis to the open field data after swim stress exposure. Also for this dataset, 2D embedding reveals that the group mean is shifted away from the control group after 45 min, but 24 hours after stress exposure the group mean overlaps again with the control group (Figure 3F,H). Next, we plotted the BFF embedding across all three experiments (Figure 3I). In order to remove potential batch effects across experiments, we normalized behavioral flow to the internal control group within each dataset (by normalizing the transition matrix of each animal to the mean of the control group, see Methods section for details). This is in notable contrast to a previous attempt, where each group was compared not to an internal control, but to all other groups combined^18^, thus potentially overestimating the power to identify effects. The resulting 2D embedding shows that yohimbine - which triggers noradrenaline release - induces very distinct changes that separate the yohimbine group very strongly from control animals. Acute stress - during which noradrenaline is released as well - shifts the animals towards the yohimbine group (45min), an effect that disappears as stress effects subside (24hrs). This shows that combining BFF with cluster stabilization allows visualizing treatment effects and comparing the impact of various manipulations across experiments and individuals in ways that were previously not possible.

### 4. Cluster transfer to new datasets and data integration

Next, we tested how well the analysis pipeline can be transferred to new datasets that were not used to perform clustering and train the clustering classifier. To this end, we performed two new experiments. In the first experiment, mice were exposed to chronic restraint stress (CRS) for 90 minutes per day on 10 consecutive days (Figure 4A), and tested in the OFT 45 minutes after the last stress exposure. In the second experiment, we triggered noradrenaline release directly in the brain using chemogenetic (DREADD, hM3Dq) activation of the locus coeruleus^26, 27^ (Figure 4B). We recorded open field behavior directly after DREADD activation. Then, we performed clustering as described above using the cluster classifier trained on the CSI, swim stress and yohimbine data (Figure 4C).

**Figure 4.**
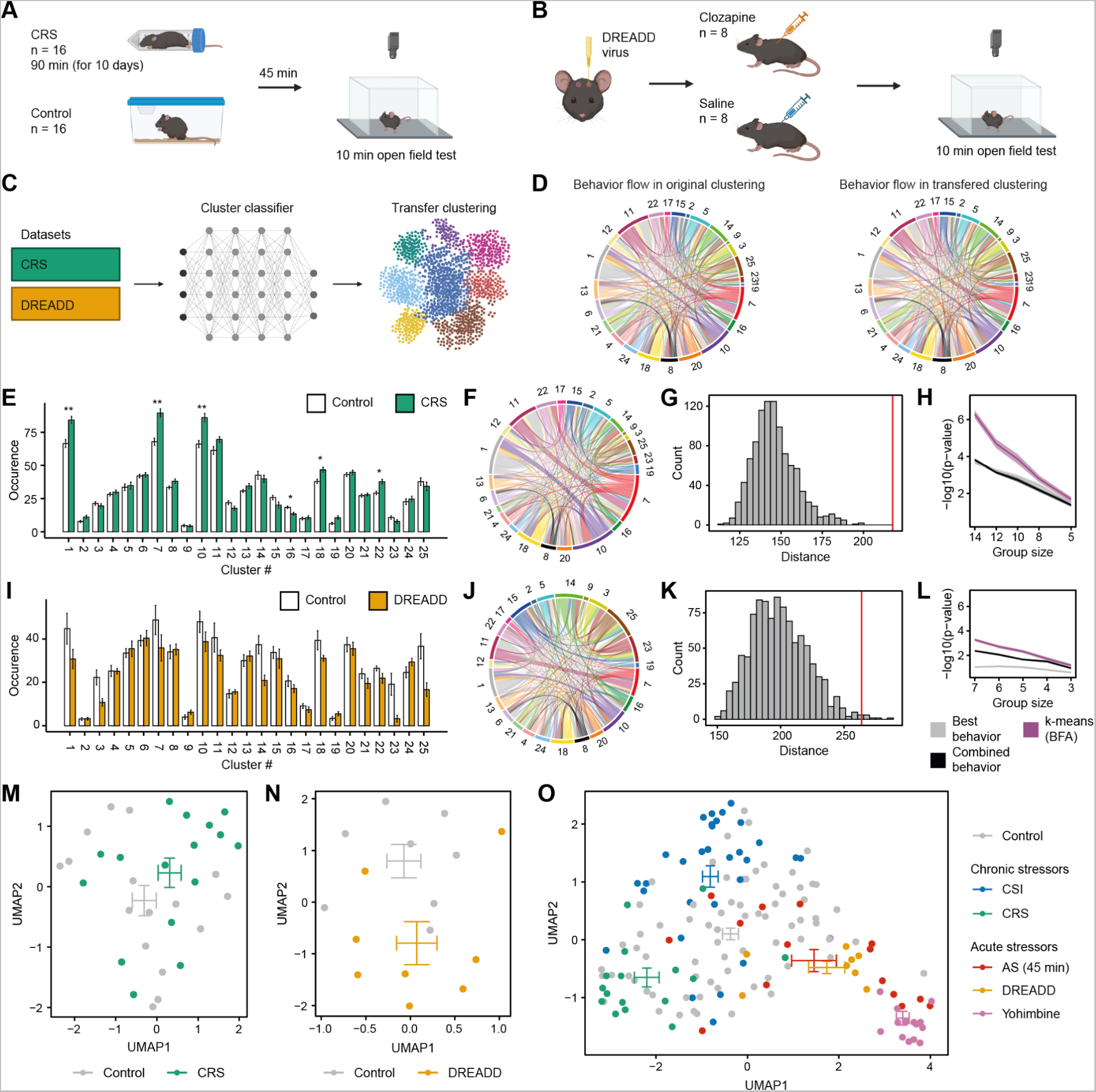
Clustering is transferable to new datasets with the same experimental setup. (A) Schematic of experimental design for OFT after chronic restraint stress (CRS), or (B) after DREADD activation of the locus coeruleus. (C) Cluster transfer to new datasets that were not used for the initial clustering. (D) Comparison of average behavior flow in control animals reveals a similar pattern between original clustering (CSI, AS and yohimbine) and transferred clustering (CRS and DREADD). (E) Quantification of cluster occurrences in CRS. (F) Absolute differences in behavioral flow in control vs. CRS. (G) BFA reveals a treatment effect for CRS (percentile=99.9, z=5.41, p<0.001). (H) Phenotype detection sensitivity in CRS. (I) Quantification of cluster occurrences after DREADD-activation of the locus coeruleus. (J) Absolute differences in behavioral flow in saline vs. clozapine. (K) BFA reveals a treatment effect for the DREADD experiment (percentile=99.2, z=2.91, p=0.002). (L) Phenotype detection sensitivity in DREADD experiment. (M) BFF using dimensionality reduction for the CRS experiment, and (N) for the DREADD experiment. Crossbars represent the average UMAP1 and UMAP2 values ± SEM for each group. (O) BFF embeddings across original (CSI, AS, yohimbine) and new (CRS, DREADD) experiments. Adj. p-values are denoted as *<0.05 and **<0.01. Error bars denote ± SEM.

The behavioral flow diagram of control animals from the two new datasets appears very similar to the behavioral flow diagram of control animals from the previous datasets (Figure 4D). Further, we observed that clusters 1 and 11 (Suppl. Video 13 and 14) mapped to onset and offset of supported rearing, respectively, consistent with the previous experiments (see Suppl. Videos 10 and 11). This demonstrates a reproducible clustering transfer for completely new datasets. Further analysis revealed 6 significant clusters in the CRS experiment (Figure 4E), but no cluster transition survived multiple testing correction (Figure 4F). BFA identified a highly significant group effect (Figure 4G) and greatly increased power to detect group differences (Figure 4H). In the DREADD experiment no significant clusters or transitions were detected (Figure 4I,J), which is likely due to the small number of animals in the control group (n=8) and variable responses between mice. However, BFA was again able to reveal a significant group effect (Figure 4K), and increased power to detect group effects compared to classical behavioral analyses (Figure 4L).

Although no cluster transitions were significantly different between groups for either experiment (Figure 4F,J), BFF clearly separated phenotypes in the 2D embedding (Figure 4M,N). We then plotted all experiments presented thus far in one 2D embedding (Figure 4O). Strikingly, all manipulations that acutely trigger noradrenaline release (acute swim, yohimbine and DREADD) were shifted away from the control groups in the same direction. In sharp contrast, the two chronic stressors (CSI and CRS) induced distinct phenotypes, with animals being shifted towards two different directions. This demonstrates that our analysis pipeline can be employed on different datasets that were not used for training the clustering classifier, and that across experiments this approach holds the potential to phenotype behavioral response profiles in ways that are consistent with the underlying brain processes.

### 5. BFF captures individual variability and allows behavioral predictions

#### Longitudinal screening of behavior

We have shown that BFF can readily resolve group differences, but even more promising is its ability to capture complex behavior on the level of individual animals. In mice exposed to acute swim stress (AS) we noted - despite the clear treatment effect - that several animals exposed to AS embedded closer to controls (Figure 4O), raising the possibility that they might have been less responsive to the effects of stress. Distinguishing responders from non-responders is one of the great challenges in preclinical research, as it would enable screening for molecular mechanisms or therapeutics, and facilitate analyses of circuit mechanisms that determine treatment success. We thus devised an experimental setup that allowed us to repeatedly screen animals in the OFT, using an inescapable footshock (IFS) paradigm, which delivers a series of strong fooshocks over 20 minutes and induces long-lasting behavioral changes in rats^28, 29^ and mice^30, 31^. Mice were tested in the OFT the day before stress exposure (OFT1), the day after stress exposure (OFT2), as well as one week afterwards (OFT3). Between OFT2 and OFT3, fear memory was assessed and mice were exposed to one extinction session per day, in which they were placed in the footshock context for 5 minutes without shock exposure (see Figure 5A). Cluster analysis revealed a strong stress-induced phenotype on OFT2, the day after footshock exposure (Figure 5B), and two behavioral transitions revealed a significant difference between groups (Figure 5C,D). These effects on cluster occurrences and transitions were not observable before footshock exposure (OFT1, Suppl. Figure 6A,B) and disappeared again after the extinction period during OFT3 (Suppl. Figure 5C,D). Next, we applied the BFA and BFF algorithms to assess performance of the animals on all three OFT tests. BFA revealed no significant difference between groups before stress exposure (OFT1), a very strong phenotype the day after stress exposure (OFT2), and again no significant group difference 7 days after stress exposure and following extinction (OFT3; Figure 5E). Then we used BFF to plot the behavioral transition dynamics in 2D together with all previous experiments, which revealed a very strong separation between groups (Figure 5F). Notably, the animals exposed to IFS clustered together with the other acute stress and noradrenaline manipulations, in line with the notion that acute stress is characterized by a strong release of noradrenaline.

**Figure 5.**
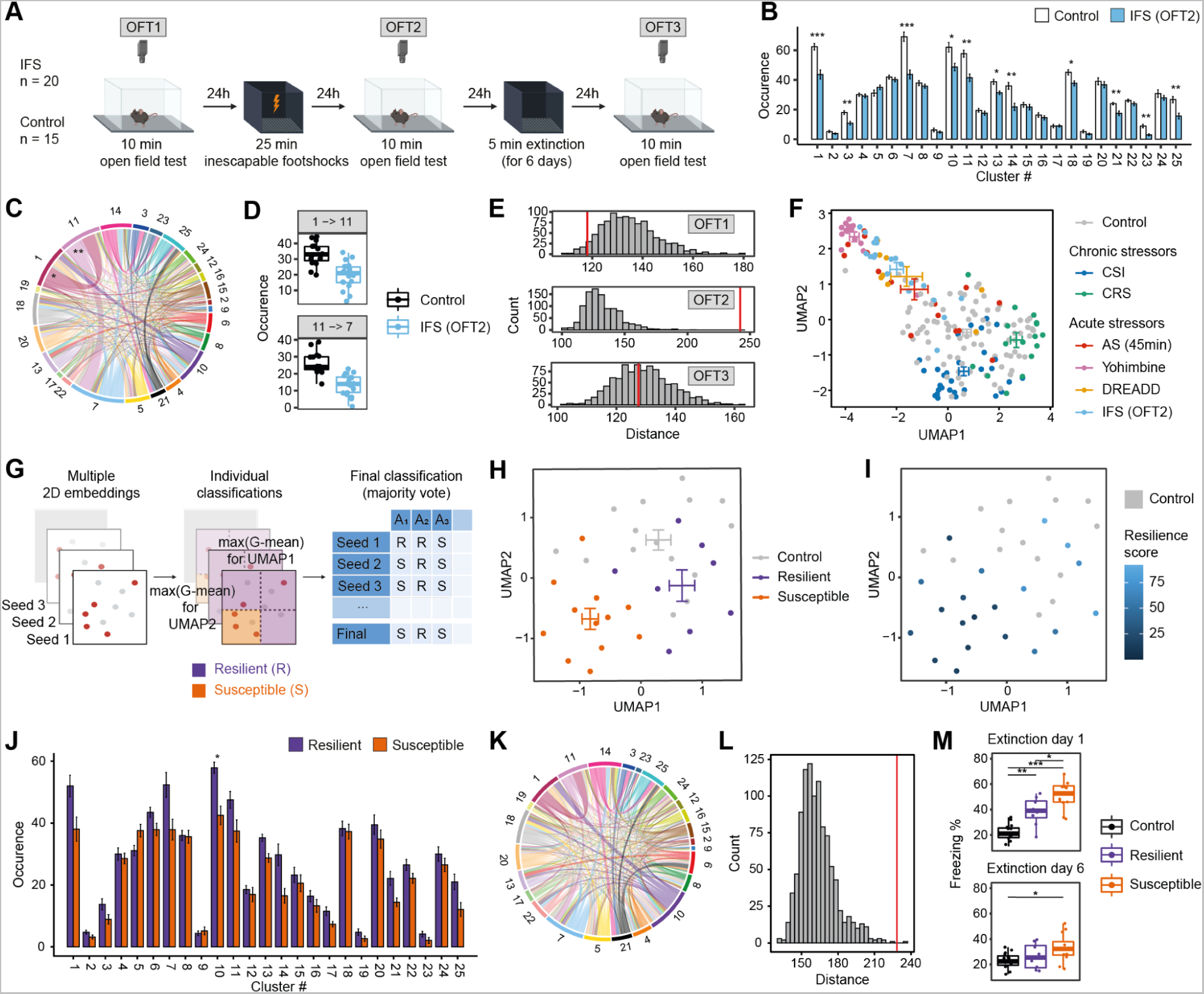
BFF captures individual variability and allows behavioral predictions. (A) Schematic of experimental design for inescapable footshock (IFS). (B) Cluster occurrences in IFS (OFT2). (C) Absolute difference in behavioral flow in control vs. IFS (OFT2). (D) Significant transition occurrences in control vs. IFS (OFT2). (E) BFA reveals a treatment effect of IFS only during OFT 2 (OFT1: percentile=3.9, z=-1.54, p=0.939; OFT2: percentile=99.9, z=7.77, p<0.001; OFT3: percentile=44.5, z=-0.18, p=0.570). (F) 2D embedding of behavioral flow across all datasets (stabilized to controls). (G) Schematic workflow of stratifying non-responding and responding groups based on multiple 2D embeddings. (H) 2D embedding of behavioral flow in IFS (OFT2) labeled for group assignment, or (I) labeled for their stress-responsivity score. (J) Cluster occurrences in non-responding vs. responding mice. (K) Absolute differences in behavioral flow in responding vs. non-responding mice. (L) BFA reveals a group effect between responding and non-responding mice (percentile=99.8, z=4.44, p<0.001). (M) Freezing response on extinction days 1 and 6 reveals differences between non-responding and responding mice (extinction day 1: F(2,32)=29.37, p<0.001; extinction day 6: F(2,32)=4.08, p=0.026). Adj. p-values are denoted as: *<0.05, **<0.01, ***<0.001. Error bars denote ± SEM.

#### Capturing stress-responsiveness in high-dimensional space

Based on the BFF results, we devised a method to determine which stress-exposed animals appear more similar to control mice (non-responders), and which mice show a stronger behavioral change in response to stress (responders). The “responder”- attribution was based on UMAP1 and UMAP2 thresholds with the highest geometric mean of recall and specificity for each dimension. To account for parameter sensitivity of the UMAP reduction, we repeated the classification into responders and non-responders for several different 2D embeddings of the same transition data by changing the random seed. The final classification was then based on a majority vote over all 2D embeddings (schematic shown in Figure 6G, see Methods for more details). This approach allows separating responders from non-responders based not on a single behavioral test, but by taking the entire high-dimensional behavioral flow data into account (Figure 5H). Beyond stratifying animals into groups, this approach allows assigning a “stress-responsivity score” to each individual mouse, based on how often the animal was classified as a non-responder in all the 2D embeddings (percentage of classifications) (Figure 6I). Using this new group assignment, we compared OFT2 performance and found - as expected - different cluster representation and behavioral flow (Figure 5J,K) and a very strong group difference using BFA (Figure 5L).

**Figure 6.**
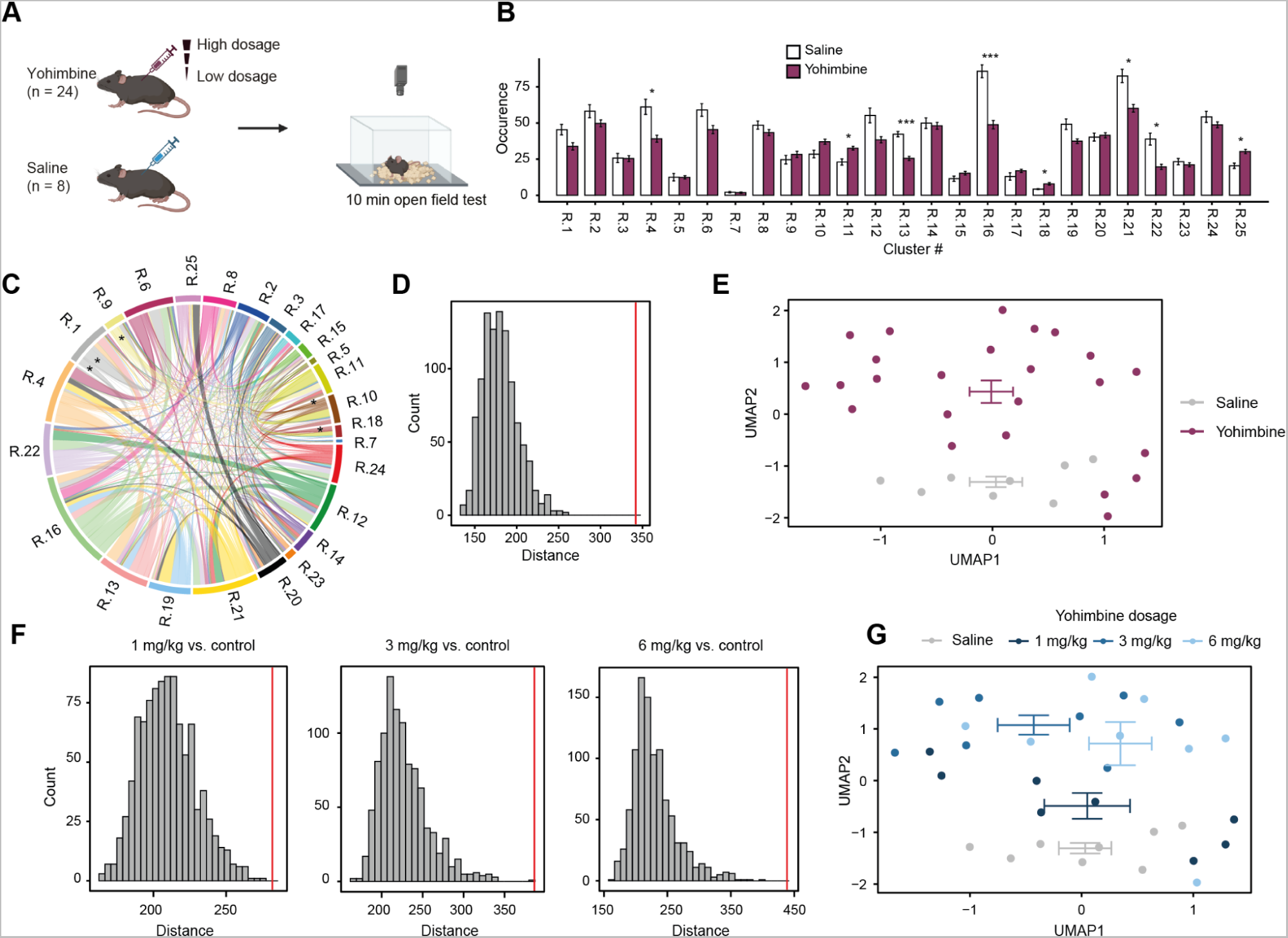
BFA and BFF are transferable to other setups. (A) Schematic showing experimental design for OFT after yohimbine or saline injection in another laboratory using a different OFT setup. These are not the same clusters as with the cluster stabilizer introduced earlier, thus they are labeled #R.1-R.25 (B) Cluster occurrences in yohimbine. (C) Absolute difference in behavioral flow in mice treated with saline vs. yohimbine. (D) BFA reveals a treatment effect for yohimbine (percentile=99.9, z=7.63, p<0.001). (E) BFF shows a separation between saline and yohimbine treatment. Crossbars represent the average UMAP1 and UMAP2 values ± SEM for each group. (F) Separate BFA of different yohimbine dosages show an increasing treatment effect (1 mg/kg: percentile=99.9, z=3.76, p<0.001; 3 mg/kg: percentile=99.8, z=5.33, p<0.001; 6 mg/kg: percentile=99.9, z=6.04, p<0.001) (G) BFF also separates the different doses of yohimbine treatment. Adj. p-values are denoted as: *<0.05, **<0.01, ***<0.001. Error bars denote ± SEM.

More interestingly, using the same group assignment we found that responders (based on OFT2 performance) showed a stronger freezing response 48 hours after prolonged footshock exposure compared to non-responders (Figure 5M), and responders also failed to fully extinguish their freezing response during 6 consecutive days of extinction training (Figure 5M, Suppl. Figure 5E).

A previous study had reported that - using a very similar experimental design in rats - the activity (distance traveled) in the open field can be used to dissociate responders from non-responders^32^. We thus used the same approach to divide mice based on ’distance traveled’ into the top and bottom 25% (Suppl. Figure 5F). However, this group assignment was not able to reveal any differences in freezing responses before or throughout extinction (Supplementary Figure 5G). This shows that BFF provides a description of stress-induced animal behaviors that is sufficiently detailed to enable a prediction about future behavioral performance on a different test.

### 6. Transferability of BFA and BFF across labs and setups

Finally, we tested whether our BFA/BFF analysis pipeline could be applied to open field data collected in an entirely different setup in another laboratory (Roche, Basel). We again devised an experiment in which mice were challenged with different doses of yohimbine. This time, mice received either vehicle injection, or injections of 1, 3 or 6 mg/kg yohimbine (Figure 6A). Five minutes after drug injection, mice were tested in the open field test for 10 minutes. Notably, the new setup was smaller (40.5cm x 40.5cm) than our original setup (45x45cm), and differed in multiple ways (see methods for details). Briefly, the floor was covered with wood chip bedding, the setup was inside a different sound attenuating enclosure, camera setup and lighting conditions were different and video acquisition was performed with 30 instead of 25 frames per second (fps). Top-view video tracking was again performed using DeepLabCut. Because our cluster classifier was not designed for stabilizing clusters across different setups and fps, we again performed k-means clustering (k=25) on these data, and then used the same analysis pipeline for BFA and BFF as described above. We find clear treatment effects using cluster occurrence (Figure 6B, note that these are not the same clusters as with the cluster stabilizer introduced earlier, thus we labeled them #R.1-R.25), transitions (Figure 6C), as well as a very strong group effect using BFA (Figure 6D). Also BFF revealed a clear separation between groups (Figure 6E).

Next, we applied BFA separately to each dose (n=8) and found that it detects strong treatment effects in all doses, with p-values getting lower as the dose gets higher (Figure 6F). Again, BFA shows much higher statistical power when compared to the quantification of cluster occurrence, which only detects a single cluster as significant at the lowest dose (1mg/kg, see Suppl. Figure 6). Finally, also BFF clearly separates the different doses of yohimbine in a 2D embedding (Figure 6E).

## Discussion

### Empowering the analysis of behavior

Big data approaches hold enormous promise, but also present major challenges. When transcriptomic screening techniques first emerged, molecular biologists swiftly embraced the vast potential of unbiased discovery, often overlooking basic statistical rigor ^33, 34^. Similar issues followed the emergence of fMRI analyses in neuroscience ^35^ and the advent of GWAS studies in medicine^36^. In each case, it took many years to recognize the substantial issue of false-positive findings stemming from inflated alpha-error probabilities. Analogously, we are in the midst of a revolution in behavioral neuroscience, where big data approaches allow us to describe and quantify rodent behavior with unprecedented resolution^4, 15^. The most advanced algorithms to assess complex behaviors quantify the total number of occurrences of each cluster or the number of transitions between clusters to identify group differences^10, 16, 18, 21, 37, 38^. However, analyzing rodent behavior is challenging, because it can be affected by multiple factors that are independent of the experimental manipulation and difficult to control. Such factors include for example the scent of the experimenter^39^, variations in maternal care^40^, hierarchies within the cage^41^ or social interactions with stressed conspecifics^42^. When this notorious variability is combined with the high number of measurements that result from data-driven approaches, stringent multiple testing correction becomes quintessential. As a result, large group numbers are required, which runs counter to the goal that these approaches set out to achieve in the first place: a more refined analysis of rodent behavior that reduces the number of animals needed for testing.

Our BFA approach addresses this issue by circumventing the multiple testing problem. Using a large number of independent behavioral datasets, we demonstrate that this approach is able to increase statistical power independent of group size, and that it is able to resolve subtle phenotypes when classic analyses fail. Further, we use a behavioral benchmarking dataset to demonstrate that BFA can be efficiently deployed using the output of several, fundamentally different clustering approaches^16, 21^. On the one hand, this suggests that the dynamic information contained within transitions between behavioral clusters - irrespective of how these clusters are identified - provides critical insights into animal behavior. On the other hand, the compatibility of our approach with other clustering algorithms allows researchers to use it as an add-on tool to determine whether a given dataset contains a treatment effect, before using more exploratory analyses to identify the nature of the behavioral differences (i.e. identify which clusters or transitions are impacted).

### BFF for deep behavioral profiling of individuals

One of the great challenges in preclinical research is to identify treatment responsiveness on the level of individual animals using behavior testing. In stress research, this typically involves determining which of the stress-exposed animals classify as resilient (they can maintain health and well-being) and which animals are susceptible (they will develop maladaptive health outcomes^43, 44^). The most popular approaches to stratify resilient/susceptible animals only use a single measure on a single test, e.g. the time they spend interacting with an unfamiliar animal after chronic social defeat^45^, the amount of sucrose consumed after stress^46^, the intensity of the startle response^47^, or the time unstressed animals explore certain areas during approach-avoidance tests^48, 49^. More elaborate strategies use several behavioral tests (e.g. to assess anxiety, social interaction, anhedonia etc.) following stress exposure, and dissociate resilient from susceptible mice by assigning composite test scores^50–52^. While the former approach is problematic because it relies on one measure on a single test, profiling animals across multiple tests is labor intensive and incompatible with some experimental designs.

As a proof-of-concept approach, we harness the ability of BFF to represent high-dimensional behavioral data collected from a single behavioral test in one “composite” datapoint per animal. We show that using BFF on open field behavior one day after a single inescapable footshock stressor can be used to predict long-term fear and fear extinction responses on the group level. This is reminiscent of recent work in humans, where complex behavioral features were collected during an approach-avoidance task to successfully predict physiological stress responses better than a clinical assessment^53^. Future work will be needed to carefully assess whether this approach can improve the predictive power compared to more elaborate screening strategies. However, the ability to behaviorally profile individual animals using a short and widely-used laboratory test, will empower biomedical research labs to integrate behavior analysis with big data approaches, such as molecular screening or high-throughput imaging.

### Limitations and outlook

Several notable limitations plague any data-driven attempt to segment behavior recordings. First, not all clusters yield behaviors that can be recognized by human observers, which limits the interpretability of findings. Second, choosing the optimal number of clusters is a well-known and difficult challenge^54^. We chose to provide a computationally efficient k-means clustering approach, based on features extracted from bodypoint-tracking with sensible temporal integration. We then quantified the observed transitions between clusters to capture the dynamic behavioral flow rather than the absolute counts of clusters. Many of the resulting clusters yield recognizable and interpretable behavioral motifs, and the observed transitions reveal meaningful behavioral sequences of mouse behavior, but many clusters (and their transitions) remain difficult to interpret. Regarding cluster numbers, many different evaluation metrics are available to determine the optimal number of clusters, and they typically do not agree with one another^15, 54^. We determined cluster numbers based on an approach that is simple and intuitive, and has been used by behavior segmentation methods before^17, 21^. However, this approach is influenced by the choice of cluster numbers used for the initial assessment (100 clusters in our case). Overall, when choosing the number of clusters, there is likely a delicate balance between signal and noise, where too few clusters result in a weak signal, and too many amplify noise. Even for a given test and setup, there is presumably no ’optimal’ number of clusters, but rather, it is contingent on the specific phenotype under investigation. Simpler phenotypes (e.g. when locomotor activity is generally affected) may be sufficiently captured with fewer clusters, while detecting more nuanced phenotypes could necessitate more clusters.

While these considerations challenge the idea of selecting a fixed number of clusters, our BFF approach showcases the advantage of defining and stabilizing a set of clusters across many experiments. By training our clustering algorithm on a subset of videos that represent a number of different experimental conditions, we stabilize the cluster assignment across experiments. We demonstrate that - within the same experimental setup - this approach can resolve behavioral phenotypes and visualize large-scale comparisons of individual animals across experimental conditions. This revealed not only a clear dissociation between acute and chronic stressors, but it identified similarities between behavioral changes induced by acute stress and noradrenergic manipulations, in line with the known biological link between acute stress and activation of the noradrenergic system through the locus coeruleus ^24, 55, 56^. Going forward, there is likely a trade-off between identifying the optimal number of clusters to maximally boost statistical power for each given experiment, and the decision to stabilize a “reasonable” number of clusters to allow direct comparisons across large numbers of datasets collected within a given laboratory or potentially across larger research consortia.

## Data availability

All video data produced in our lab (411 separate recordings), corresponding pose estimation data and metadata has been deposited online and can be accessed under https://zenodo.org/record/8186065. All video data and pose estimation data produced by Roche Pharma (32 separate recordings) can be accessed under https://zenodo.org/record/8188683. The BehaviorFlow package and furthermore any script used to analyze our data and generate the manuscript figures can be accessed under (https://github.com/ETHZ-INS/BehaviorFlow)

## Acknowledgments

The lab of J.B. is supported by the, ETH Zurich, ETH Project Grant ETH-20 19-1, SNSF Grants 310030_172889 and 310030_204372, the Botnar Research Centre for Child Health, the Swiss 3R Competence Center, Roche, and the Hochschulmedizin Zürich Flagship project STRESS.

We thank the staff of the EPIC for the excellent animal care and their service to our animal facility. We thank Julia Bode for maintaining the animal colony, Runzhong (Yvonne) Zhang for practical experimental help, Sebastian Leimbacher for contributions to code development and behavior testing, and Pierre-Luc Germain for critical reading of the manuscript and for supporting our data infrastructure. We thank Brigitte Algeyer for conducting the experiment at Roche and Yan-Ping Zhang, Damian Roqueiro and Francesca Tozzi for comments on the manuscript. The scope of Roche funding did not extend to the acute swim stress experiments, and swim stress was employed here as a stressor, not to measure despair as conducted in the forced swim test.

## Author Contributions

LvZ conceived experiments, developed algorithms, conducted data analyses, produced figures, interpreted results, wrote manuscript. FR developed algorithms, conducted data analyses, produced figures, wrote manuscript. OS conducted experiments (CSI, AS, yohimbine). RW conducted experiments (CRS, IFS). MP conducted experiment (AS). SD conducted experiment (LC-DREADD). EC Supervised experiments (yohimbine) and provided resources. JB conceived experiments, interpreted results, provided resources and funding, wrote manuscript.

## Methods

### Animals

Mice were maintained in a temperature- and humidity-controlled facility on a 12 h reversed light– dark cycle (lights off at 08:15 a.m.) with food and water ad libitum. Mice were housed in groups of 5 per-cage and used for experiments when 2.5–4 months old unless stated otherwise. For each experiment, mice of the same age were used in all experimental groups to rule out confounding effects of age. All tests were conducted during the animals’ active (dark) phase from 10 a.m. to 6 p.m. Mice were habituated to the colony room for at least 2 weeks before experimentation. Mice were single housed 24h before behavioral testing in order to standardize their environment and avoid disturbing cagemates during testing^57, 58^. All procedures were carried out in accordance to Swiss cantonal regulations for animal experimentation and were approved under licenses: ZH155/2015, ZH161/2017, ZH106/2020, ZH067/2022.

### Open field test (OFT)

Open field testing took place inside sound insulated, ventilated multi conditioning chambers (TSE Systems Ltd, Germany). The open field arena (45 × 45 × 40 cm [L × W × H]) consisted of four transparent Plexiglas walls and a light gray PVC floor. Animals were tested for 10 min under dim lighting (4 lux). Animals were removed from their home cage and placed directly into the center of the open field. The doors of the conditioning chamber were then swiftly closed. Video recording at 25 fps was triggered by infra-red beam break upon the mouse entering the arena. Pose estimation was performed using DeepLabCut and data were processed using custom R Scripts that are available online (https://github.com/ETHZ-INS/DLCAnalyzer) to measure distance, time in center, supported rears and unsupported rears^9^.

### Chronic social instability (CSI)

The CSI procedure was carried out as previously described^59^ on male C57BL/6J (C57BL/6JRj) mice (n=59) obtained from Janvier (France). The mice arrived at the lab aged between postnatal day 21–23 housed in groups of 5. Upon arrival they were ear-tagged and split randomly (by cage) into either the CSI or control group. The CSI mice underwent the CSI paradigm, which consisted of briefly placing all CSI mice (n = 30) into a larger cage, from which they were then randomly assigned to new cages. Mice in the control group were similarly handled, however, they entered the larger cage only with their cagemates before they were all returned to their original cage. The mice were subjected to these cage changes twice a week (Tuesday/Friday) for seven weeks, during the last cage change the mice were returned to their original cage and allowed to rest for 5 weeks prior to any further testing.

### Acute swim (AS)

Male (C57BL/6JRj) mice (n=30) were obtained from Janvier (France). Acute swim (AS) mice (n=15) had to swim in a plastic beaker (20 cm diameter, 25 cm deep) filled to 17 cm with 17.9–18.1 °C water for 6 minutes before they were placed back into their home cages. Control mice (n=15) remained in their homecage until further testing in the Open field. Open field testing was performed 45 min and 24 hours after the stress.

### Yohimbine injections

Male C57BL/6J (C57BL/6JRj) mice (n=20) were obtained from Janvier (France). Before open field testing mice were hand-restrained and received one of 15 dosages of yohimbine ranging from 0.4 mg/kg - 6 mg/kg (n=15) or vehicle (saline 0.9%) (n=5) (i.p.). Mice were immediately placed into the center of the open field test arena following their restraint and i.p. injection.

### Chronic restraint stress (CRS)

C57BL/6J (C57BL/6JRj) male (n=16) and female (n=16) mice were obtained from Janvier (France) and housed in groups of 4 animals per cage. Upon arrival, animals were randomly split into a control and chronic restraint stress (CRS) group by cage. CRS animals were placed in a 50 ml Falcon tube with a large air hole for 90 minutes for 10 consecutive days, while control animals were gently handled daily. On day 10, CRS animals were placed in the open field test arena 45 minutes after the end of the restraint stress.

### Chemogenetic activation of locus coeruleus (DREADD)

Heterozygous C57BL/6-Tg(Dbh-icre)1Gsc female (n=16) mice were subjected to stereotactic brain injections, as described previously^27^. The mice were anesthetized with isoflurane and placed in a stereotaxic frame. For analgesia, animals received a subcutaneous injection of 2 mg/kg Meloxicam and a local anesthetic (Emla cream; 5% lidocaine, 5% prilocaine) before and after surgery. A pneumatic injector (Narishige, IM-11-2) and calibrated microcapillaries (Sigma-Aldrich, P0549) were used to inject 1 μL of virus (ssAAV-5/2-hSyn1-dlox-hM3D(Gq)_mCherry(rev)-dlox-WPRE-hGHp(A); physical titer: 4 x 10E12 vg/ml) bilaterally into the locus coeruleus (coordinates from bregma: anterior/posterior −5.4 mm, medial/lateral ± 1.0 mm, dorsal/ventral −3.8 mm). 3 weeks after surgery, animals were tested in the open field arena for 20 min. For consistency in this study, we only used the first 10 min of recordings. Animals were injected i.p. with 0.03 mg/kg clozapine (Sigma-Aldrich, Steinheim, Germany) (n=8) or saline 0.9% (n=8) and placed directly into the center of the open field test arena.

### Yohimbine experiment from Roche

Male C57BL/6J mice (n=32) were obtained from Charles River Laboratories (Saint Germain sur l’Arbresle, France) and single housed in GM500 cages (Tecniplast) upon arrival in the test facility (Roche Innovation Center, Basel) to prevent aggression. Cages were supplemented with nesting material and two pieces of enrichment that were changed during each cage change. Mice were given *ad libitum* access to food (Standard Diet; Kliba Nafag) and water, and temperature and humidity were continuously monitored and controlled to 22°C ± 2°C and 50 ± 10 %, respectively. The holding and test room were maintained on a 12h:12h light: dark cycle, with lights transitioning to fully on by 06:00. Ethical approval for this study was provided by the Federal Food Safety and Veterinary Office of Switzerland. All animal experiments were conducted in strict adherence to the Swiss federal ordinance on animal protection and welfare as well as according to the rules of the Association for Assessment and Accreditation of Laboratory Animal Care International (AAALAC), and with the explicit approval of the local veterinary authorities (License BS2448).

Mice were acclimatized to the facility for 1 week prior to the locomotor activity test. On the day of the test, mice (n=8 per treatment group) were randomly allocated to receive either Yohimbine at 1, 3 or 6 mg/kg (i.p.) or its vehicle (0.3% Tween 80 in 0.9% Saline; i.p.), 5 minutes prior to being placed into the center of the test arena. The dose volume was 10 mL/Kg and mice weighed on average 24.5g (min / max.: 23g / 27.4g) at the time of the test. The test arena was a clear Perspex chamber (41 x 41 x 30.5 cm), held within a sound- and light-attenuating cubicle (Omnitech Electronics, USA), and retrofitted with a camera positioned above the chamber to enable continuous video capture at 30 fps of locomotor activity during the 30-minute test. For behavioral flow analysis, only the first 10 minutes of recording were used.

### Inescapable footshock (IFS)

Male C57BL/6J (C57BL/6JRj) mice (n=35) were obtained from Janvier (France). Upon arrival, animals were randomly split into a control and inescapable footshock (IFS) group and single-housed one day prior to behavioral testing. All animals were first tested in the open field test for 10 min (OFT1). One day later, IFS animals were placed inside the TSE multi conditioning systems’ black fear conditioning arena. After 5 min of rest, the animals received 19 foot shocks (0.5 sec, 1 mA) over 20 min. Shocks were randomly distributed over the 20 minutes to be 30, 60 or 90 seconds apart. Control animals were placed in the same boxes without receiving a footshock. 24 hours later, all animals were placed in the open field test for 10 min (OFT2). All animals were placed in the black fear conditioning arena for 5 minutes daily for 6 consecutive days one day before a final open field test (OFT3).

### Pose estimation and tracking based open field analysis

DeepLabCut 2.0.7 was used to track 13 body points and the 4 corners of the open field test arena. Tracked points included nose, headcentre, neck, right ear (earr), left ear (earl), bodycentre, bodycentre left (bcl), bodycentre right (bcr), left hip (hipl), right hip (hipr), tailbase, tail center, tail tip. In addition the 4 corners of the open field were tracked to automatically detect the open field arena boundaries in each recording. The networks for different tests were trained using 10–20 frames from multiple randomly selected videos for 250,000–1,030,000 iterations. X and Y coordinates of DLC-tracking data were imported into R Studio (v 3.6.1) and processed with the DLCAnalyzer package^9^. Points relating to the arenas were used to define the arenas *in silico* by using their median XY coordinates. Values of points with low likelihood (<0.95) and points tracked outside an existence polygon (arena scaled by a factor 1.3) were removed and interpolated using the R package “imputeTS” (v.3.2). The speed and acceleration of each point was determined by integrating the animal’s position over time. The pixel-to-cm conversion ratio for each video was determined by comparing the volume of the arena *in silico* in px^2^ to the measured size of the arena in cm^2^. Zones of interest were calculated from the arena using polygon-modification functions . Furthermore, we applied a previously trained supervised classifier^9^ to quantify supported and unsupported rears on a per frame basis.

### Statistical analysis of open field data

In order to assess group differences based on tracking data prior to clustering, we used automated metrics from the DLCAnalyzer package^9^ such as distance moved, time spent in center and number of supported and unsupported rears. Significance was assessed with a standard parametric t-test.

### Generation of feature data for k-means clustering

We used the pose estimation data (*N* ∼ 15000 frames) to generate a set of *m* = 41 features (see table below) resulting in a large numeric matrix **X_i_** ∈ ℝ*^N x m^* for each recording (here denoted with index *i*). 5 different types of features were used: acceleration of points, distance between point-pairs, angle between two point-pairs, distance of point to closest border, and area of a polygon spanned by multiple points (Table 1).

**Table 1:**
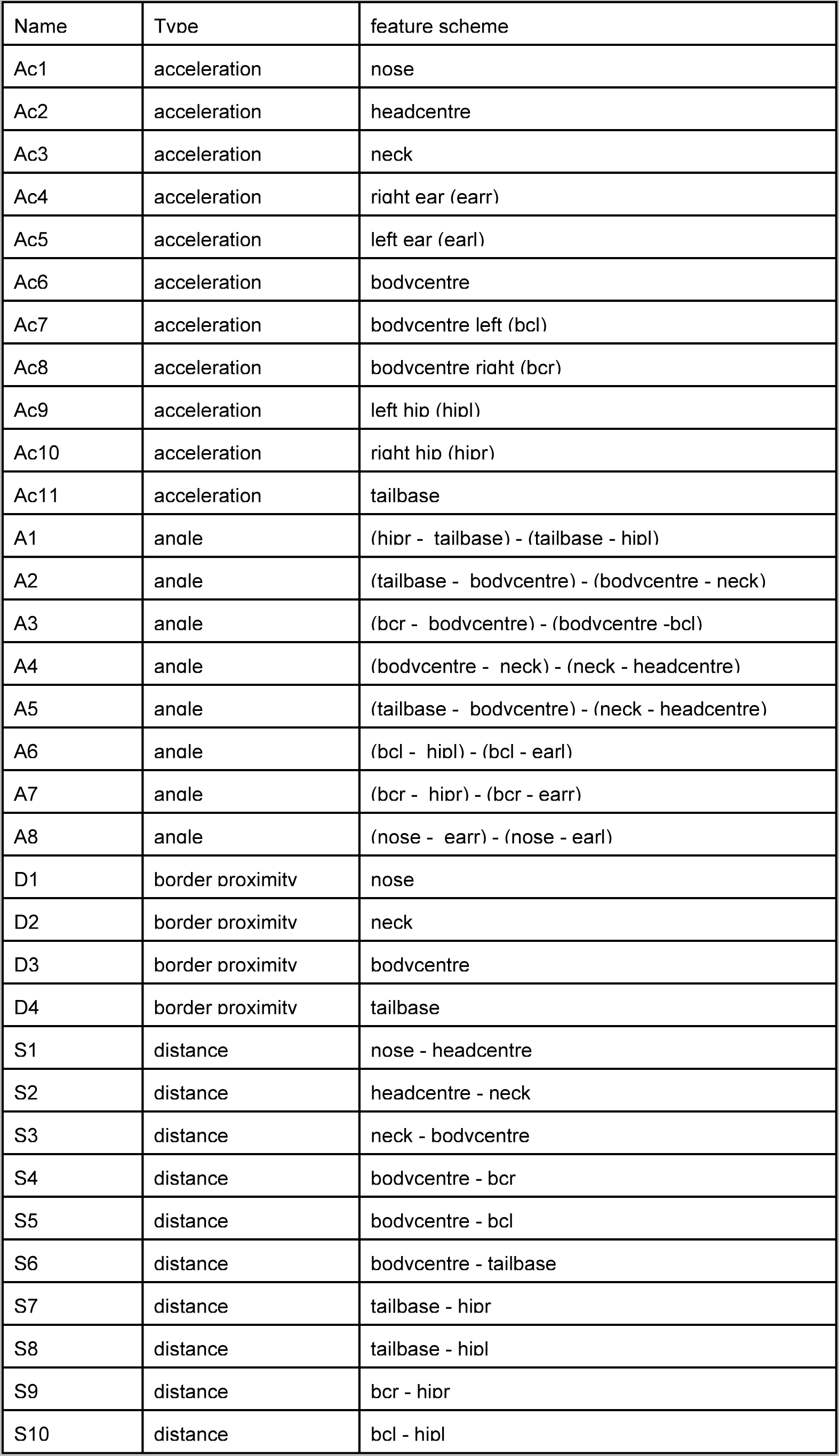
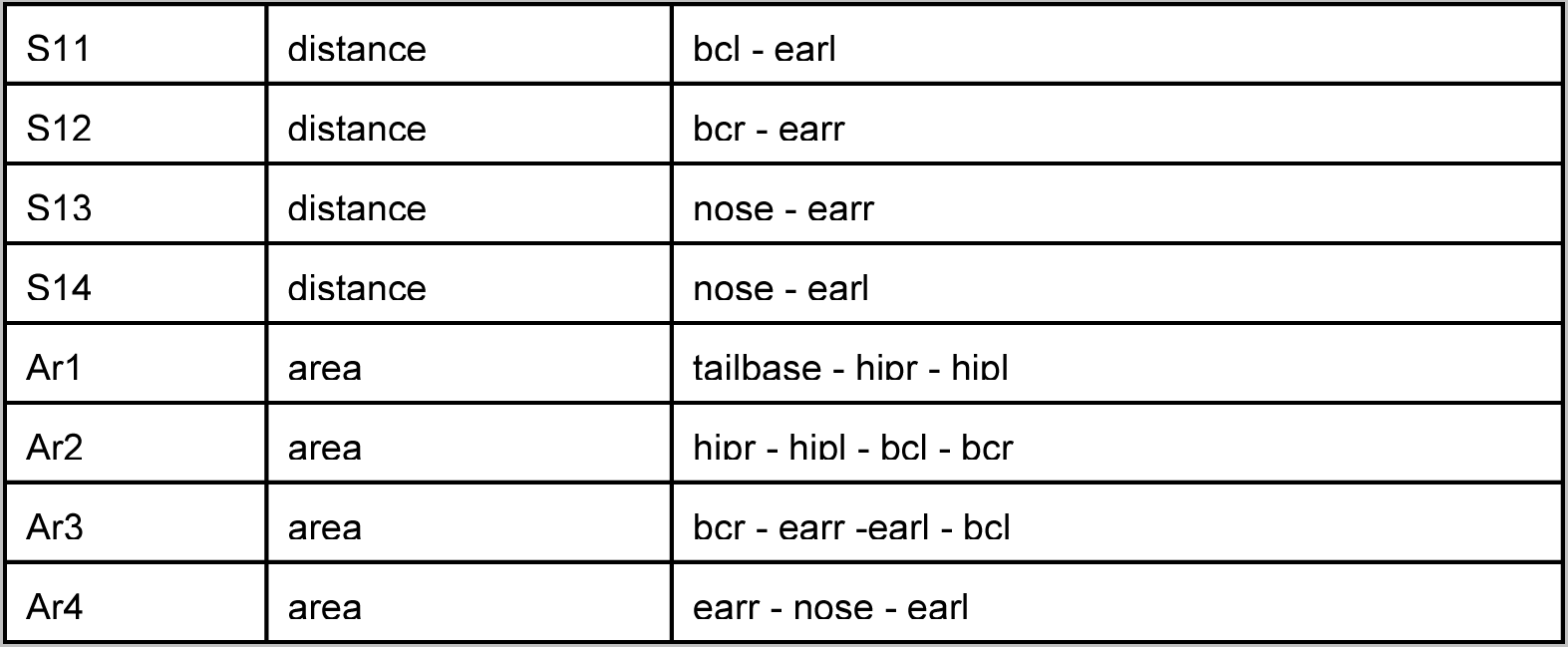
Feature data.

| Name | Type | feature scheme |
| --- | --- | --- |
| Ac1 | acceleration | nose |
| Ac2 | acceleration | headcentre |
| Ac3 | acceleration | neck |
| Ac4 | acceleration | right ear (earr) |
| Ac5 | acceleration | left ear (earl) |
| Ac6 | acceleration | bodycentre |
| Ac7 | acceleration | bodycentre left (bcl) |
| Ac8 | acceleration | bodycentre right (bcr) |
| Ac9 | acceleration | left hip (hipl) |
| Ac10 | acceleration | right hip (hipr) |
| Ac11 | acceleration | tailbase |
| A1 | angle | (hipr - tailbase) - (tailbase - hipl) |
| A2 | angle | (tailbase - bodycentre) - (bodycentre - neck) |
| A3 | angle | (bcr - bodycentre) - (bodycentre - bcl) |
| A4 | angle | (bodycentre - neck) - (neck - headcentre) |
| A5 | angle | (tailbase - bodycentre) - (neck - headcentre) |
| A6 | angle | (bcl - hipl) - (bcl - earl) |
| A7 | angle | (bcr - hipr) - (bcr - earr) |
| A8 | angle | (nose - earr) - (nose - earl) |
| D1 | border proximity | nose |
| D2 | border proximity | neck |
| D3 | border proximity | bodycentre |
| D4 | border proximity | tailbase |
| S1 | distance | nose - headcentre |
| S2 | distance | headcentre - neck |
| S3 | distance | neck - bodycentre |
| S4 | distance | bodycentre - bcr |
| S5 | distance | bodycentre - bcl |
| S6 | distance | bodycentre - tailbase |
| S7 | distance | tailbase - hipr |
| S8 | distance | tailbase - hipl |
| S9 | distance | bcr - hipr |
| S10 | distance | bcl - hipl |
| S11 | distance | bcl - earl |
| S12 | distance | bcr - earr |
| S13 | distance | nose - earr |
| S14 | distance | nose - earl |
| Ar1 | area | tailbase - hipr - hipl |
| Ar2 | area | hipr - hipl - bcl - bcr |
| Ar3 | area | bcr - earr - earl - bcl |
| Ar4 | area | earr - nose - earl |

Feature data was then first normalized on a per recording level. Z-score normalization was used for distances and area features, angle data (in rad) was not normalized and border proximities and accelerations were scaled linearly (with a factor of 0.1 and 4 respectively). These feature data were furthermore expanded over sequences of +- 15 frames (*t* = 31) centered on each frame to generate a larger feature set of *m * t* = 1271 values for each frame resulting in **X_temporal,i_** ∈ ℝ*^N x (m * t)^*.

### Determining best number of clusters

For determining the best number of clusters, we applied an approach described previously by others^17, 21^. We first ran the clustering for a total number of 100 clusters. For each cluster, we then computed the proportion of image frames assigned to it. To determine the best number of clusters, we added the cluster proportions (sorted from high to low proportion) and chose the number of clusters which contain 95% image frames as the best one.

### k-means clustering

For k-means clustering, we selected a random subset of 20 samples per experiment from the chronic social instability, the acute swim stress and the yohimbine injection experiments (*s* = 60). All feature data of each frame for these subsets were combined into one single large matrix **X_clustering_** ∈ ℝ*^(N * s) x (m * t)^* that was then z-score normalized across columns. The normalized feature matrix was k-means clustered using the function bigkmeans() of the R package “biganalytics” (v.1.1.21)

### B-SOiD

As a comparison to k-means clustering, we ran B-SOiD^16^ on the chronic social instability dataset. We followed the steps described on the tutorial webpage (bsoid.org). In short, we trained B-SOiD on a random subsample of 20 files containing the pose estimation computed by DLC. Due to computational issues, we used a reduced set of 9 tracking points including nose, headcentre, neck, bodycentre, bodycentre left (bcl), bodycentre right (bcr), left hip (hipl), right hip (hipr), and tailbase as input for B-SOiD. We explored different “minimum cluster size ranges” and ended up with a range between 0.10 and 0.95. The remaining 39 files (not used for clustering) got their clusters assigned using a trained random forest classifier.

### VAME

We ran VAME^21^(v.1.1) using PyTorch(v.1.7.0) and followed the workflow as described in the publication. We ran VAME on all 59 pose estimation files from DLC. The 11 tracking points including nose, headcentre, neck, right ear (earr), left ear (earl), bodycentre, bodycentre left (bcl), bodycentre right (bcr), left hip (hipl), right hip (hipr), and tailbase used as input were egocentric aligned to the two tracking points nose and tailbase. Before clustering, we changed the parameters “n_cluster” to 80 (see above how we determined the best number of clusters), “pose confidence” to 0.95 and “n_features” to 22 in the configuration file.

### Clustering classifier

To transfer clustering to larger or new datasets, we trained a sequential neural network to imitate the clustering results obtained with k-means. We used the framework designed in a previous publication^9^ with the clusters obtained from k-means as ground truth labeling data and **X_clustering_** as input data. We used R packages for tensorflow (v. 2.9.0) and keras (v. 2.8.0) to design and train the neural network. We used a neural network with a single hidden layer of 1024 units (using “relu” activation) and an input shape of 1271 followed by a dropout layer with a rate of 0.4 to prevent overfitting. We used an output layer with 25 output neurons using the “softmax” activation function. Data was shuffled before training for 30 epochs with a batch size of 512. We used the “categorical crossentropy” loss function, “rmsprop” as optimizer and “accuracy” as metric during training. Clustering classifiers were then applied to **X_temporal_** of each individual recording to obtain the final clustering results.

### Clustering classifier assessment

To assess the performance of the clustering classifiers, we performed a 10 fold cross-validation. All 60 recordings in the clustering (= training) set were randomly shuffled before sequentially a different set of 6 recordings were set aside for validation each time and the training was performed on the remaining 54 recordings only. Then, for each cross-validation pass we calculated precision, recall and F1 score on a per cluster basis.

### Label data processing

We used the newly written BehaviorFlow Package to process all label data (= cluster assigned to each frame) from the k-means classifier, VAME and B-SOiD. To remove noise and single frame misclassifications, we processed this data by first smoothing all labels using a sliding window of +- 5 frames and selecting the most abundant categorical value across the window using the SmoothLabels_US() function. Next, we calculated metrics such as number of clusters, behavior onset/offsets and time spent in each cluster on a per recording basis using the CalculateMetrics() function. We then calculated the transition matrix across all label groups for each recording independently using the AddTransitionMatrixData(). This function first removes any repeating labels from each label vector to create an occurrence vector. Then, this occurrence vector and the same vector shifted by 1 element are used to calculate the contingency table using the table() function of the base package of R resulting in a transition matrix **T_i_** ∈ W *^NC^ ^x^ ^NC^* where i denotes the i-th recording and NC the number of clusters. To calculate stabilized transition matrices **T_stabilized,i_** ∈ ℝ *^NC^ ^x^ ^NC^* we used the function CalculateStabilizedTransitions() for defined subsets and control recordings. The stabilized transition matrix is defined as the difference to the mean of the control recordings **(1)**. Next, we calculated the confusion matrix across all label groups using the AddConfusionMatrix() function. This function calculates a contingency table for a source-target pair using the table() function of the base package of R with the full label vector of the source and the full label vector of the target label group.

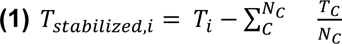

*T_i_* = Transition matrix of recording i, *N_C_* = number of control group recordings, *T_C_* = Transitionmatrix of control group recording C

### Statistical analysis of labeling data (two group analysis)

We used the TwoGroupAnalysis() function of the BehaviorFlow Package for the statistical analysis of label data. This function runs a number of statistical tests to test for group differences on the cluster usage level and the transition level on each label group. For number of cluster occurrences and time spent with clusters, a simple parametric t-test followed by a Benjamini-Yekutieli multiple testing correction (using the function t.test() and p.adjust() of R) was used. The same test was also applied to individual transitions. To test for overall differences across all transitions (referred to as behavioral flow analysis or BFA), we first calculated the group-wise mean transition matrix and then the absolute Manhattan distance between the two groups based on these mean matrices **(2)**. We then used a bootstrapping approach to estimate a null distribution of the inter group distance from random groupings. We randomly shuffled the group assignment vector 1000 times and calculated the inter group distance for each sampling. We then used this to calculate the percentile for non-parametric statistics **(3)**. Next, we calculated mean and standard deviation from the null distribution and z **(4)** for parametric tests. We used the error function to calculate the parametric right-tailed p-value from z using the function erf() from the R package “pracma” (v.2.3.8) for the hypothesis true distance > null distribution distances **(5)**.

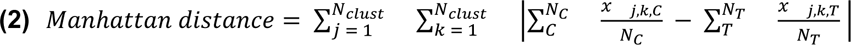

*N_clust_* = number of clusters, *N_C_* = number of control group recordings, *N_T_* = number of test group recordings, *x_i,j,r_* = number of transitions from cluster *i* to cluster *j* in recording *i*

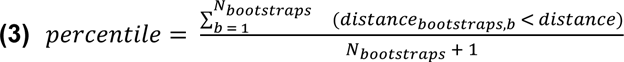

*N_bootstraps_* = number of bootstraps, *distance_bootstraps,b_* = Manhattan distance from b-th bootstrapping, *distance* = true group Manhattan distance from **(2)**

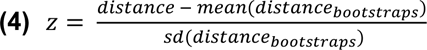

*distance* = true group Manhattan distance from **(2),** *distance_bootstraps_* = Manhattan distances obtained from bootstrapping, *mean()* = arithmetic mean, *sd() =* standard deviation

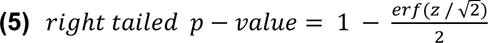

*z* from **(4),** *erf() =* error function

### *in silico* sensitivity assay

To create the sensitivity curves, we designed an *in silico* assay that step wise reduces group sizes and randomly selects a subset of both groups (ensuring that they are equally sized) prior to running a two group comparison. For the CSI dataset we used group sizes 25, 20,15, 10, and 5; for the AS dataset we used group sizes of 13, 11, 9, 7, and 5; for the CRS dataset we used group sizes of 14, 12, 10, 8, and 5 and for the DREADD dataset group sizes of 7, 6, 5, 4, and 3. To better estimate the p-value at each step, we performed 50 random samplings followed by a two group comparison each time. For “best behavior”, we picked the lowest p-value from the 4 classical readouts (distance, time spent in center, number of supported and unsupported rears) for each random sampling and group size. For “combined behavior”, we devised an analysis method that leverages all automated readouts from the DLCAnalyzer package. Due to the highly correlated nature of some readouts (i.e raw distance and distance moving), we first applied a principal component analysis on all readouts. Then, we selected the principal components that cumulatively explain at least 90% of the variance and fitted a linear model with the formula *group ∼ PC1 + … + PCN*, (where “group” is a binary grouping variable, PC1 is the first principal component and PCN the Nth principal component) using the lm() method of the R. This model was then tested against a null model using an analysis of variance method to assess if a group difference can be verified based on the principal component data. Last, we −log10 transformed all p-values prior to calculating mean and standard deviation of the p-values for each dataset.

### Log-linear modeling

To test for an association between yohimbine dosage and the occurrence of transitions between clusters, we used a log-linear model. We first transformed both the yohimbine dosages and the transition occurrences using the natural logarithm. We then fitted a linear model to these two variables using the lm() method from R.

### 2D embedding

For 2D embedding, we used the function Plot2DEmbedding() of the BehaviorFlow Package. As input data, the stabilized transition matrix (**T_stabilized,i_**) was used for each recording. We then used a UMAP embedding with the function umap() from the R package “M3C” (v.1.16.0). Points were colored by relevant groups and +- standard error of the mean (SEM) ranges (based on UMAP1 and UMAP2 coordinates) were added to the plot for visual aid.

### Grouping of resilient and susceptible animals

To determine resilient and susceptible groups of animals based on 2D embeddings of transition data, first calculated UMAP1 and UMAP2 coordinates using the Plot2DEmbedding() function of the BehaviorFlow Package. Then, we separately assessed for their classification performance (stressed vs. control) using receiver operating characteristic (ROC) curves (roc() from R package “pROC”, v.1.18.0). The best performing thresholds in distinguishing groups using either UMAP1 or UMAP2 were determined using the highest geometric mean (G-mean) value computed between recall and specificity. The two thresholds then divide the 2D embedding into quadrants, and the one with the highest number of stressed animals was used to define them as susceptible while stressed animals in the other three quadrants were defined as resilient. To adjust for the sensitivity of the umap() function to setting a random seed, we repeated this procedure 101 times for different random seeds and let a majority vote (>50%) over all classifications decide on the final group assignment for each stressed animal.

## References

1. Skinner, B. F. The behavior of organisms: an experimental analysis. 457, (1938).

2. Darwin, C. & Prodger, P. The Expression of the Emotions in Man and Animals. (Oxford University Press, 1998).

3. Kalueff, A. V. & Zimbardo, P. G. Behavioral neuroscience, exploration, and K.C. Montgomery’s legacy. Brain Res. Rev. 53, 328–331 (2007).

4. von Ziegler, L., Sturman, O. & Bohacek, J. Big behavior: challenges and opportunities in a new era of deep behavior profiling. Neuropsychopharmacology 46, 33–44 (2021).

5. Segalin, C. et al. The Mouse Action Recognition System (MARS) software pipeline for automated analysis of social behaviors in mice. Elife 10, (2021).

6. Nath, T., et al. Using DeepLabCut for 3D markerless pose estimation across species and behaviors. http://biorxiv.org/lookup/doi/10.1101/476531 (2018).

7. Lauer, J. et al. Multi-animal pose estimation, identification and tracking with DeepLabCut. Nat. Methods 19, 496–504 (2022).

8. Pereira, T. D. et al. SLEAP: A deep learning system for multi-animal pose tracking. Nat. Methods 19, 486–495 (2022).

9. Sturman, O. et al. Deep learning-based behavioral analysis reaches human accuracy and is capable of outperforming commercial solutions. Neuropsychopharmacology 45, 1942–1952 (2020).

10. Marks, M. et al. Deep-learning-based identification, tracking, pose estimation and behaviour classification of interacting primates and mice in complex environments. Nat Mach Intell 4, 331–340 (2022).

11. Nilsson, S. R. O., et al. Simple Behavioral Analysis (SimBA) – an open source toolkit for computer classification of complex social behaviors in experimental animals. bioRxiv 2020.04.19.049452 (2020) doi:10.1101/2020.04.19.049452.

12. Nourizonoz, A. et al. EthoLoop: automated closed-loop neuroethology in naturalistic environments. Nat. Methods 17, 1052–1059 (2020).

13. Bohnslav, J. P. et al. DeepEthogram, a machine learning pipeline for supervised behavior classification from raw pixels. Elife 10, (2021).

14. Datta, S. R., Anderson, D. J., Branson, K., Perona, P. & Leifer, A. Computational Neuroethology: A Call to Action. Neuron 104, 11–24 (2019).

15. Pereira, T. D., Shaevitz, J. W. & Murthy, M. Quantifying behavior to understand the brain. Nat. Neurosci. 23, 1537–1549 (2020).

16. Hsu, A. I. & Yttri, E. A. B-SOiD, an open-source unsupervised algorithm for identification and fast prediction of behaviors. Nat. Commun. 12, 5188 (2021).

17. Wiltschko, A. B. et al. Mapping Sub-Second Structure in Mouse Behavior. Neuron 88, 1121–1135 (12/2015).

18. Wiltschko, A. B. et al. Revealing the structure of pharmacobehavioral space through motion sequencing. Nat. Neurosci. 23, 1433–1443 (2020).

19. Bordes, J. et al. Automatically annotated motion tracking identifies a distinct social behavioral profile following chronic social defeat stress. Nat. Commun. 14, 4319 (2023).

20. Berman, G. J., Choi, D. M., Bialek, W. & Shaevitz, J. W. Mapping the stereotyped behaviour of freely moving fruit flies. J. R. Soc. Interface 11, (2014).

21. Luxem, K. et al. Identifying behavioral structure from deep variational embeddings of animal motion. Commun Biol 5, 1–15 (2022).

22. Sturman, O. et al. Chronic adolescent stress increases exploratory behavior but does not appear to change the acute stress response in adult male C57BL/6 mice. Neurobiol Stress 15, 100388 (2021).

23. Kagiampaki, Z. et al. Sensitive multicolor indicators for monitoring norepinephrine in vivo. Nat. Methods 1–11 (2023).

24. Privitera, M., et al. Noradrenaline release from the locus coeruleus shapes stress-induced hippocampal gene expression. bioRxiv 2023.02.02.526661 (2023) doi:10.1101/2023.02.02.526661.

25. von Ziegler, L. M. et al. Multiomic profiling of the acute stress response in the mouse hippocampus. Nat. Commun. 13, 1824 (2022).

26. Privitera, M. et al. A complete pupillometry toolbox for real-time monitoring of locus coeruleus activity in rodents. Nat. Protoc. 15, 2301–2320 (2020).

27. Zerbi, V. et al. Rapid Reconfiguration of the Functional Connectome after Chemogenetic Locus Coeruleus Activation. Neuron 103, 702–718.e5 (2019).

28. Musazzi, L., Tornese, P., Sala, N. & Popoli, M. Acute stress is not acute: sustained enhancement of glutamate release after acute stress involves readily releasable pool size and synapsin I activation. Mol. Psychiatry 22, 1226–1227 (9/2017).

29. Vollmayr, B. & Henn, F. A. Learned helplessness in the rat: improvements in validity and reliability. Brain Res. Brain Res. Protoc. 8, 1–7 (2001).

30. Lecca, S. et al. Rescue of GABAB and GIRK function in the lateral habenula by protein phosphatase 2A inhibition ameliorates depression-like phenotypes in mice. Nat. Med. 22, 254–261 (2016).

31. Nuno-Perez, A. et al. Stress undermines reward-guided cognitive performance through synaptic depression in the lateral habenula. Neuron 109, 947–956.e5 (2021).

32. Colucci, P. et al. Predicting susceptibility and resilience in an animal model of post-traumatic stress disorder (PTSD). Transl. Psychiatry 10, 243 (2020).

33. Pounds, S. & Morris, S. W. Estimating the occurrence of false positives and false negatives in microarray studies by approximating and partitioning the empirical distribution of p-values. Bioinformatics 19, 1236–1242 (2003).

34. Storey, J. D. & Tibshirani, R. Statistical significance for genomewide studies. Proc. Natl. Acad. Sci. U. S. A. 100, 9440–9445 (2003).

35. Eklund, A., Nichols, T. E. & Knutsson, H. Cluster failure: Why fMRI inferences for spatial extent have inflated false-positive rates. Proceedings of the National Academy of Sciences 113, 7900–7905 (2016).

36. Sesia, M., Bates, S., Candès, E., Marchini, J. & Sabatti, C. False discovery rate control in genome-wide association studies with population structure. Proceedings of the National Academy of Sciences 118, e2105841118 (2021).

37. Huang, K. et al. A hierarchical 3D-motion learning framework for animal spontaneous behavior mapping. Nat. Commun. 12, 2784 (2021).

38. Jia, Y. et al. Selfee, self-supervised features extraction of animal behaviors. Elife 11, (2022).

39. Georgiou, P. et al. Experimenters’ sex modulates mouse behaviors and neural responses to ketamine via corticotropin releasing factor. Nat. Neurosci. 25, 1191–1200 (2022).

40. Weaver, I. C. G., Meaney, M. J. & Szyf, M. Maternal care effects on the hippocampal transcriptome and anxiety-mediated behaviors in the offspring that are reversible in adulthood. Proc. Natl. Acad. Sci. U. S. A. 103, 3480–3485 (2006).

41. LeClair, K. B. et al. Individual history of winning and hierarchy landscape influence stress susceptibility in mice. Elife 10, (2021).

42. Sterley, T.-L. et al. Social transmission and buffering of synaptic changes after stress. Nat. Neurosci. 21, 393–403 (2018).

43. Holly, E. N. & Miczek, K. A. Capturing Individual Differences: Challenges in Animal Models of Posttraumatic Stress Disorder and Drug Abuse. Biological psychiatry vol. 78 816–818 (2015).

44. de Boer, S. F., Buwalda, B. & Koolhaas, J. M. Untangling the neurobiology of coping styles in rodents: Towards neural mechanisms underlying individual differences in disease susceptibility. Neurosci. Biobehav. Rev. 74, 401–422 (2017).

45. Krishnan, V. et al. Molecular adaptations underlying susceptibility and resistance to social defeat in brain reward regions. Cell 131, 391–404 (2007).

46. Tornese, P. et al. Chronic mild stress induces anhedonic behavior and changes in glutamate release, BDNF trafficking and dendrite morphology only in stress vulnerable rats. The rapid restorative action of ketamine. Neurobiol Stress 10, 100160 (2019).

47. Torrisi, S. A. et al. A novel arousal-based individual screening reveals susceptibility and resilience to PTSD-like phenotypes in mice. Neurobiol Stress 14, 100286 (2021).

48. Hollis, F. et al. Mitochondrial function in the brain links anxiety with social subordination. Proc. Natl. Acad. Sci. U. S. A. 112, 15486–15491 (2015).

49. Nasca, C. et al. Multidimensional Predictors of Susceptibility and Resilience to Social Defeat Stress. Biol. Psychiatry 86, 483–491 (2019).

50. Ardi, Z., Albrecht, A., Richter-Levin, A., Saha, R. & Richter-Levin, G. Behavioral profiling as a translational approach in an animal model of posttraumatic stress disorder. Neurobiol. Dis. 88, 139–147 (04/2016).

51. Ritov, G., Boltyansky, B. & Richter-Levin, G. A novel approach to PTSD modeling in rats reveals alternating patterns of limbic activity in different types of stress reaction. Mol. Psychiatry 21, 630–641 (5/2016).

52. Dirven, B. C. J. et al. Longitudinal assessment of amygdala activity in mice susceptible to trauma. Psychoneuroendocrinology 145, 105912 (2022).

53. Rodrigues, J., Studer, E., Streuber, S., Meyer, N. & Sandi, C. Locomotion in virtual environments predicts cardiovascular responsiveness to subsequent stressful challenges. Nat. Commun. 11, 5904 (2020).

54. Ikotun, A. M., Ezugwu, A. E., Abualigah, L., Abuhaija, B. & Heming, J. K-means clustering algorithms: A comprehensive review, variants analysis, and advances in the era of big data. Inf. Sci. 622, 178–210 (2023).

55. McCall, J. G. et al. CRH Engagement of the Locus Coeruleus Noradrenergic System Mediates Stress-Induced Anxiety. Neuron 87, 605–620 (2015).

56. Itoi, K. & Sugimoto, N. The Brainstem Noradrenergic Systems in Stress, Anxiety and Depression. J. Neuroendocrinol. 22, 355–361 (05/2010).

57. Bohacek, J., Manuella, F., Roszkowski, M. & Mansuy, I. M. Hippocampal gene expression induced by cold swim stress depends on sex and handling. Psychoneuroendocrinology 52, 1–12 (2015).

58. Roszkowski, M. et al. Rapid stress-induced transcriptomic changes in the brain depend on beta-adrenergic signaling. Neuropharmacology 107, 329–338 (2016).

59. Schmidt, M. V. et al. A novel chronic social stress paradigm in female mice. Horm. Behav. 57, 415–420 (4/2010)

