## Supplementary Figures 1-6 for "Analysis of behavioral flow resolves latent phenotypes"

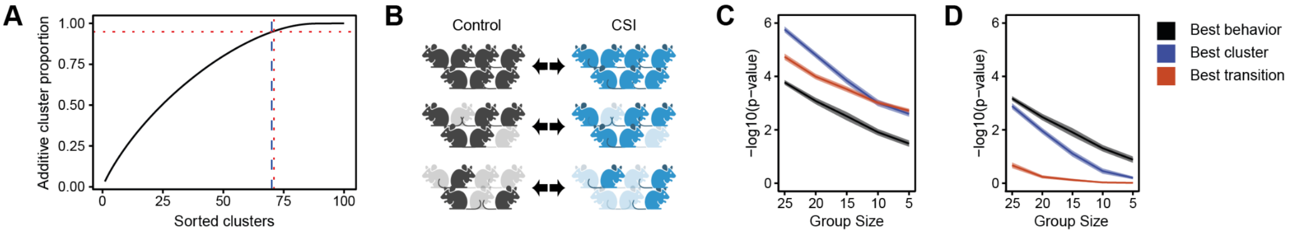


**Supplementary Figure 1.** (A) Determining the optimal number of clusters for k-means. Vertical red-dashed line marks the number of clusters (=71) which represent 95% of all frames (horizontal red-dashed line), and the blue dashed line marks the number of clusters (=70) we used for the CSI analysis. (B) Schematic of *in silico* approach to generate random subsets of each group of mice to run multiple two-group comparisons while gradually reducing group sizes. (C) Phenotype detection sensitivity in CSI with unadjusted p-values. (D) Sensitivity in CSI with adjusted p-values appropriately applying multiple testing correction.

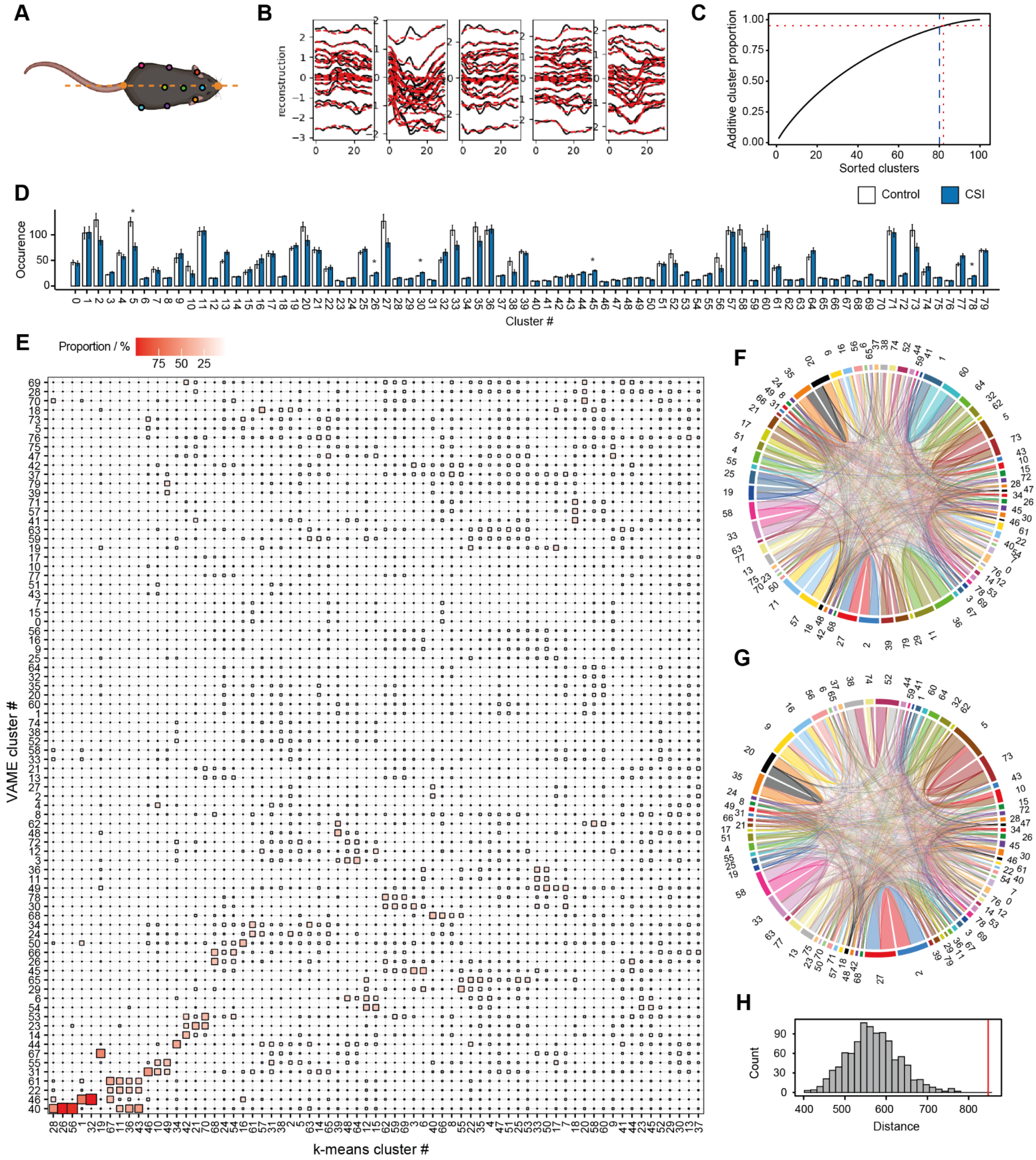


**Supplementary Figure 2.** (A) Tracking points used for VAME with egocentric alignment (orange dashed line). (B) Reconstruction of tracking points by VAME (over 30 frames). (C) Determining the optimal number of clusters for VAME. Vertical red-dashed line marks the number of clusters (=82) which represent 95% of all frames (horizontal red-dashed line), and the blue dashed line marks the number of clusters (=80) we used for the CSI analysis. (D) VAME cluster occurrences in CSI. (E) Mapping k-means clusters to VAME clusters. (F) Average behavioral flow over all animals. (G) Absolute difference in behavioral flow in CSI vs. control. For each cluster, the absolute difference in the observed number of transitions between groups is plotted. (H) BFA result for CSI using transitions between VAME clusters (percentile=99.9, z=4.42, p<0.001). Adj. p-values are denoted as *<0.05, error bars denote ± SEM.

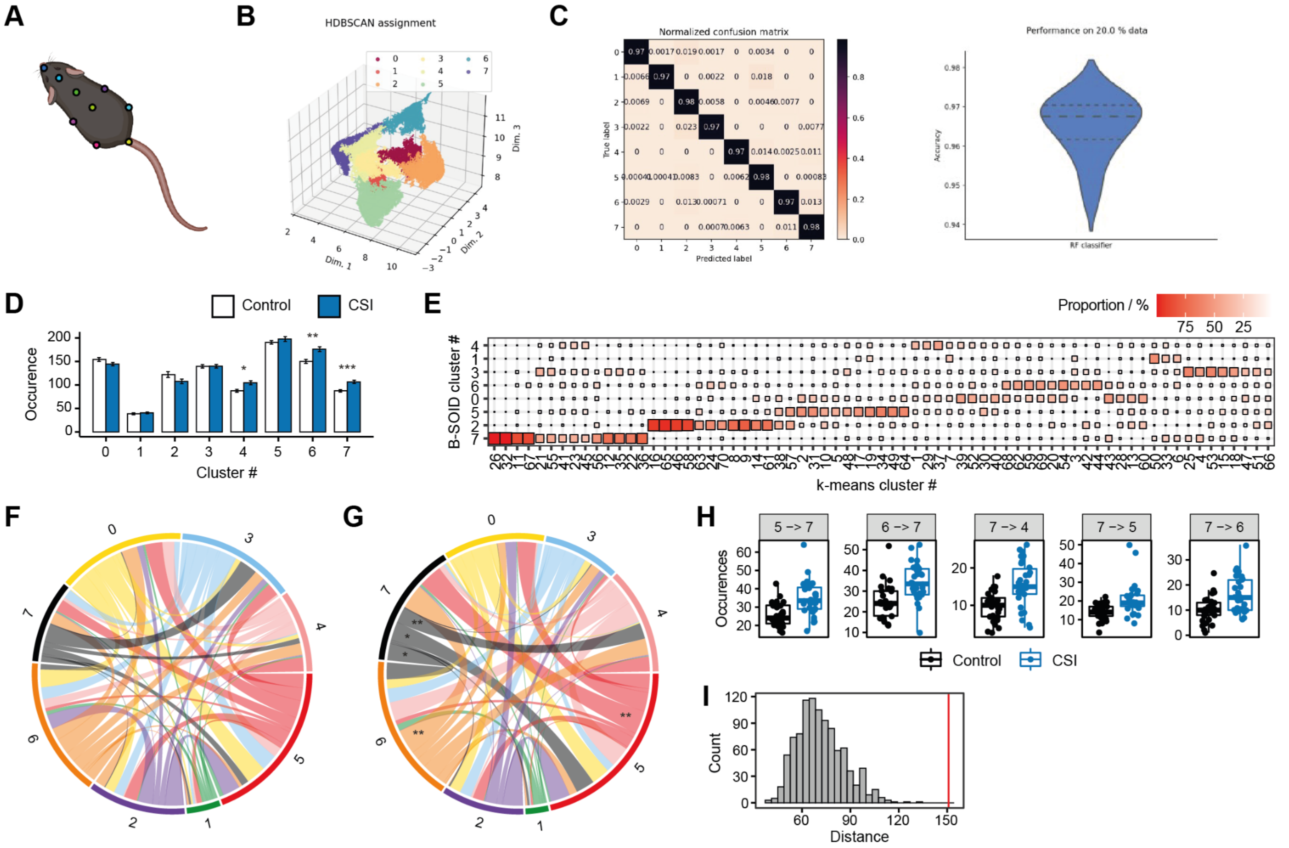

**Supplementary Figure 3.** (A) Tracking points used for B-SOiD. (B) UMAP visualization of cluster assignment in B-SOiD. (C) Performance of random forest classifier on training (left) and test (right) data. (D) B-SOiD cluster occurrences in CSI. (E) Mapping k-means clusters to B-SOiD clusters. (F) Average behavioral flow over all animals. (G) Absolute difference in behavioral flow in CSI vs. control. For each cluster, the absolute difference in the observed number of transitions between groups is plotted. (H) Transitions that are significantly different (adj. $p<0.05$) between CSI and controls. (I) BFA result for CSI using transitions between B-SOiD clusters (percentile=99.9, $z=5.52$, $p<0.001$). Adj. p-values are denoted as: *<0.05, **<0.01, ***<0.001. Error bars denote ± SEM.

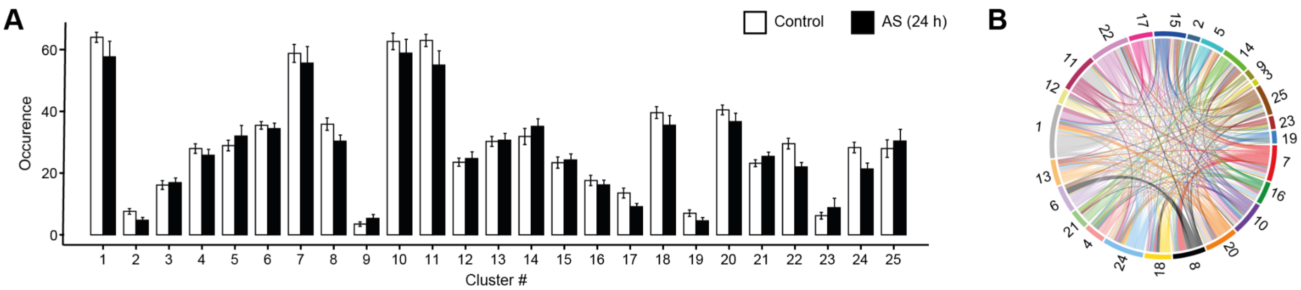


**Supplementary Figure 4.** (A) Cluster occurrences in AS (24 h). (B) Absolute difference of behavioral flow in control vs. AS (24 h). Error bars denote ± SEM.

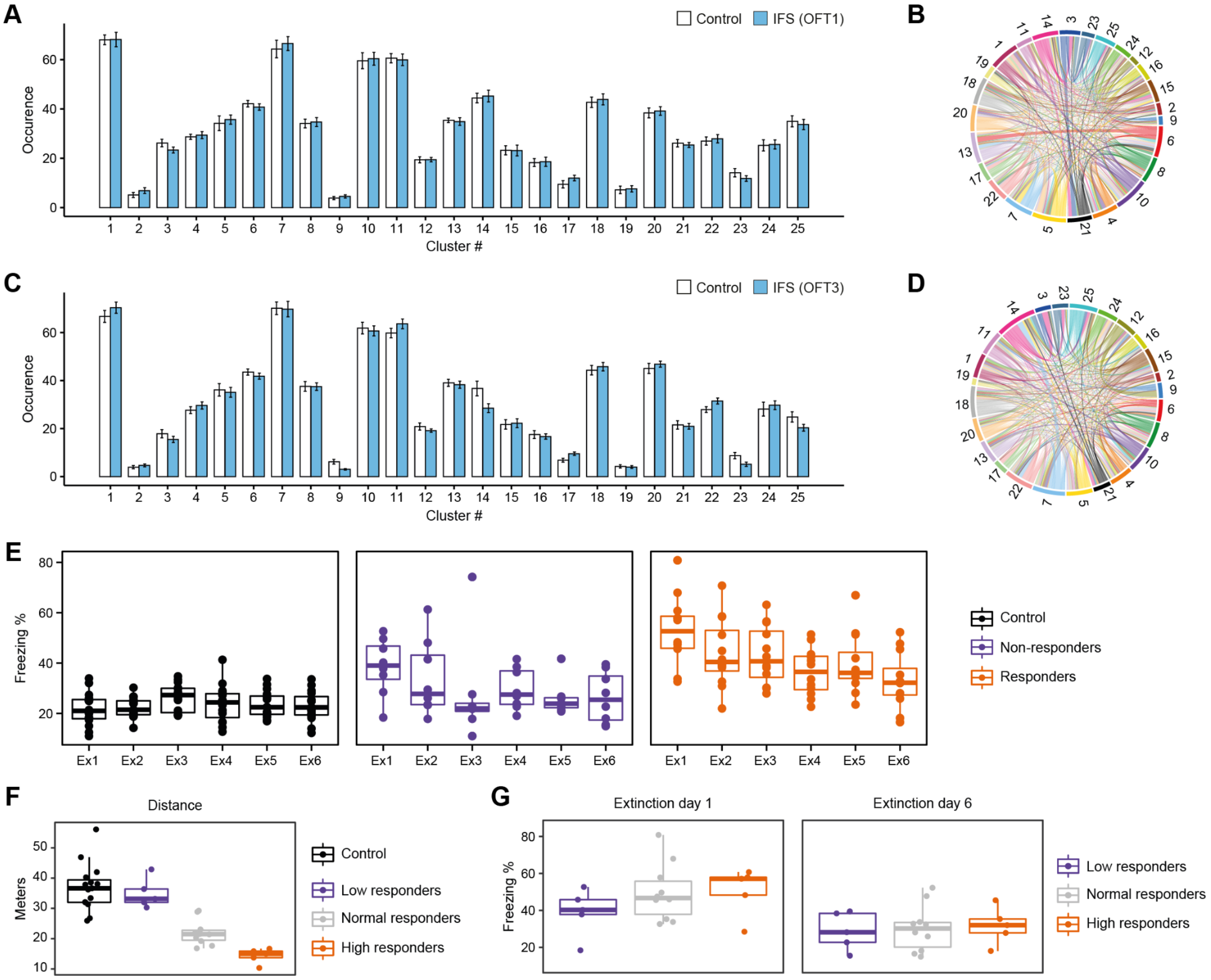

**Supplementary Figure 5.** (A) Cluster occurrences in IFS (OFT1). (B) Absolute difference in behavioral flow in control vs. IFS (OFT1). (C) Cluster occurrences in IFS (OFT3). (D) Absolute difference in behavioral flow in control vs. IFS (OFT3). (E) Freezing response across the extinction period for control, non-responder and responder mice. (F) Grouping of IFS mice into low, normal and high responders based on distance. (G) Freezing response on extinction days 1 and 6 reveals no differences between low, normal and high responding mice (extinction day 1: $F(2,17)=1.10$, $p=0.355$; extinction day 6: $F(2,17)=0.08$, $p=0.924$). Error bars denote ± SEM.

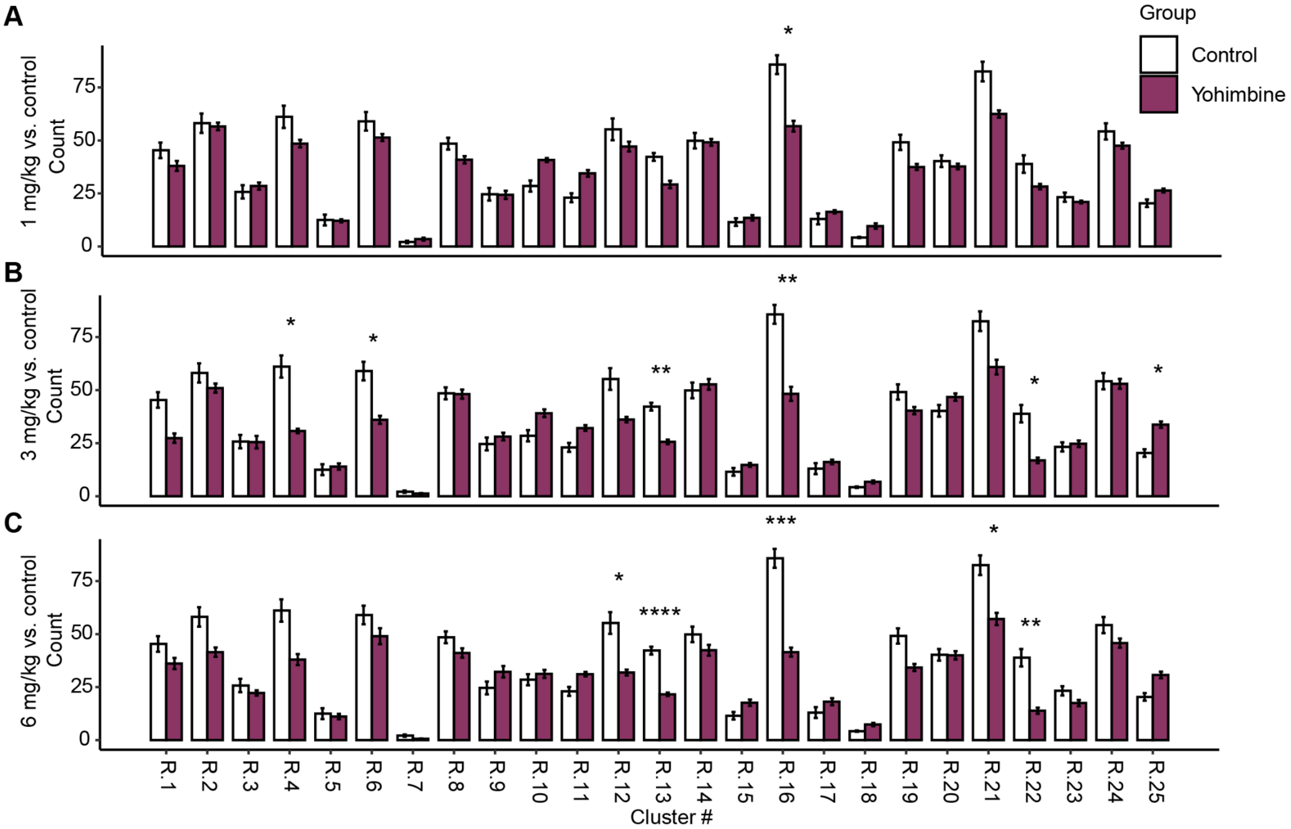


**Supplementary Figure 6.** (A) Cluster occurrences in 1mg/kg yohimbine vs. control (B) Cluster occurrences in 3mg/kg yohimbine vs. control (C) Cluster occurrences in 6mg/kg yohimbine vs. control. Adj. p-values are denoted as: *<0.05, **<0.01, ***<0.001, ****<0.0001. Error bars denote ± SEM.
